# Reverse engineering of a mechanistic model of gene expression using metastability and temporal dynamics

**DOI:** 10.1101/2021.06.01.446414

**Authors:** Elias Ventre

**Affiliations:** ENS de Lyon, CNRS UMR 5239, Laboratory of Biology and Modelling of the Cell, Lyon, France; Inria Center Grenoble Rhone-Alpes, Equipe Dracula, Villeurbanne, France; Univ Lyon, Université Claude Bernard Lyon 1, CNRS UMR 5208, Institut Camille Jordan, Villeurbanne, France

**Keywords:** single cell, gene regulation network, inference, metastability, transcriptional bursting, machine learning

## Abstract

Differentiation can be modeled at the single cell level as a stochastic process resulting from the dynamical functioning of an underlying Gene Regulatory Network (GRN), driving stem or progenitor cells to one or many differentiated cell types. Metastability seems inherent to differentiation process as a consequence of the limited number of cell types. Moreover, mRNA is known to be generally produced by bursts, which can give rise to highly variable non-Gaussian behavior, making the estimation of a GRN from transcriptional profiles challenging. In this article, we present CARDAMOM (Cell type Analysis from scRna-seq Data achieved from a Mixture MOdel), a new algorithm for inferring a GRN from timestamped scRNA-seq data, which crucially exploits these notions of metastability and transcriptional bursting. We show that such inference can be seen as the successive resolution of as many regression problem as timepoints, after a preliminary clustering of the whole set of cells with regards to their associated bursts frequency. We demonstrate the ability of CARDAMOM to infer a reliable GRN from in silico expression datasets, with good computational speed. To the best of our knowledge, this is the first description of a method which uses the concept of metastability for performing GRN inference.

## Introduction

Differentiation is the process whereby a cell acquires a specific phenotype, by differential gene expression as a function of time. Measuring how gene expression changes as differentiation proceeds is therefore of essence to understand differentiation. Advances in measurement technologies now allow to obtain gene expression levels at the single cell level. It offers a much more accurate view than population-based measurements, that has been obscured by mean population-based averaging [25], [9]. It has among other things established that there is a high cell-to-cell variability in gene expression, and that this variability has to be taken into account when examining a differentiation process at the single-cell level [30], [3], [46], [28], [41], [33], [13], [47].

A popular vision of the cellular evolution during differentiation, introduced by Waddington in [52], is to compare cells to marbles following probabilistic trajectories, as they roll through a developmental landscape of ridges and valleys. The landscape is generally described by the graph of a potential function *V* : Ω → ℝ, where Ω denotes the space where these trajectories evolve, and is called the gene expression space. In that space, a cell can be described by a vector, where each coordinate represents the expression products of a gene [17], [31].

A cell has theoretically as many states as the combination of proteins and mRNAs quantity possibly associated to each gene, which is potentially huge [7]. But metastability, that is the coexistence of multiple stable states, seems inherent to cell differentiation processes as evidenced by limited number of existing cellular phenotypes [32], [4]. Since [20] and [18], many authors have identified cell types with the basins of attraction of a dynamical system modeling the differentiation process. In that context, noise in gene expression, or cell to cell heterogeneity, appears to be closely related to the transition between these cell types [12]. This provides a rationale for coarse-graining stochastic models of gene expression into reduced processes on a limited number of metastable basins, seen as cell types. Such reduction has been studied mostly in the context of stochastic diffusion [54], [53], [57], but also for stochastic hybrid systems [23]. This last case is particularly interesting because the basins can be classified regarding to the modes associated to the jumps frequency of the discrete variable, leading to local approximations of the potential function *V* describing the landscape [51].

This landscape is often regarded to be shaped by an underlying gene regulatory network (GRN), which appears as a powerful abstraction for describing interactions between genes through their proteins production. The construction of GRNs from literature being a very time-consuming and labor intensive process, and sometimes impossible due to the limitation of our current knowledge, their automated reconstruction from large datasets has become a classic task in systems biology [44]. This task is notoriously difficult, in particular when dealing with single-cell transcriptomics, the bursty synthesis of mRNAs [58], [35] giving rise to highly variable and non-Gaussian expression data [26]. The methods that are used cover a wide range of statistical and modeling tools [1], including the analysis of stochastic models of gene expression [16], [5]. In the latter case, expression datasets are identified to independent samples of the time-varying distribution describing the process. GRN inference can then be seen as the reconstruction of the most-likely GRN from a set of partial observations of independent realizations of the model.

To our knowledge, the use of metastability for performing such reverse engineering has not been studied yet. The main contribution of this article is to derive an efficient algorithm for linking the landscape analysis of a mechanistic model of gene expression using metastability, to the most-likely associated GRN parameters. In Section 1, we are going to present a mechanistic model of gene expression developed in [16], which describes single-cell dynamics associated to a GRN with transcriptional bursting. With a methodology similar to the one developed in [51], we perform its reduction into a discrete coarse-grained model on a limited number of metastable basins. We deduce from this reduction an approximation of the time-dependent proteins distribution of the original model, presented in Section 2, which appears as a mixture of Gamma distributions. We develop in Section 3 a statistical method for linking the parameters of such mixture to the GRN parameters of the model. In Section 4, we extend this method for estimating GRN parameters from scRNA-seq timestamped datasets. We show in Section 5 the accuracy of the method for *in silico* datasets simulated from the mechanistic model with various networks of different sizes. Finally, we discuss more precisely in Section 6 the interpretation of the method in terms of landscape, its applicability to real datasets, and we highlight its similarity with a machine learning approach.

We draw the reader’s attention to the fact that the problem of inferring a GRN using a mechanistic model with transcriptional bursting has been recently elsewhere using distinct mathematical tools [15]. We are currently setting up a collaborative effort for benchmarking both algorithms on in silico generated data as well as on real datasets from the literature. This benchmarking is therefore beyond the scope of the present article and will be exposed elsewhere.

## 1 GRN model and reduction

### 1.1 Mechanistic model

The model which is used throughout this article has been introduced in [16]. It is based on a hybrid version of the well-established two-state model of gene expression [21], [38], where a gene is described by the state of a promoter, which can be {*on, off*}. If the promoter is *on*, mRNAs are transcribed at a rate *s*_0_, which are then translated into proteins at a rate *s*_1_. Degradation of both mRNAs and proteins occurs respectively at a rate *d*_0_ and *d*_1_. The transitions between the states *on* and *off* occur at exponential times of rates *k*_*on*_ and *k*_*off*_. We consider the so-called bursty regime of this model, when *k*_*on*_ « *k*_*off*_, which corresponds to the experimentally observed situation where active periods are short but characterised by a high transcription rate, thereby generating bursts of mRNA [34], [43], [50]. We describe the random times at which these bursts occur by an exponential law of parameter *k*_*on*_, and their random intensity by an exponential law of parameter *k*_*off*_ */s*_0_ (see Figure 1). This model is compatible with real single-cell data, as mRNAs quantity at the steady state follows a Gamma distribution, which is known to describe accurately continuous single-cell data [2].

**Figure 1:**
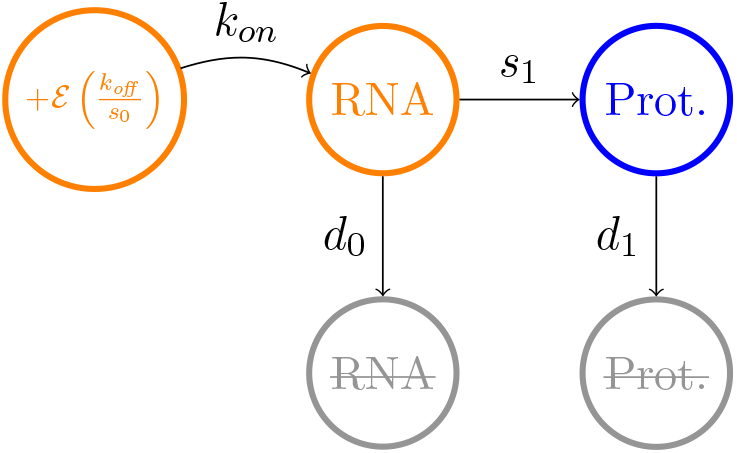
Approximation of the two-states model of gene expression in the bursty regime.

Neglecting the molecular noise associated to mRNA and protein quantities, we obtain the following mathematical description of the model:

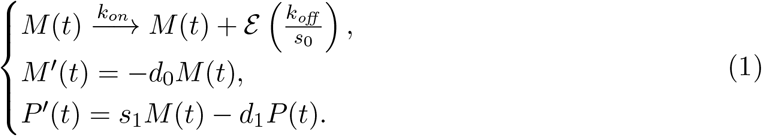

where *M* (*t*) and *P* (*t*) denote respectively the mRNA and protein concentration at time *t* and 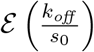 is an exponential law of mean 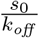. The key idea for studying a GRN is to embed this model into a network. Denoting the number of genes by *n*, the vector (*M, P*) describing the process is then of dimension 2*n*. The burst rates for each gene *i* are characterized by two gene-specific functions 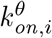 and *k*_*off* ,*i*_. For the sake of simplicity, we consider that *k*_*off* ,*i*_ does not depend on the protein level (*i*.*e* that proteins do not affect the quantity of mRNAs which are transcribed during a burst). To take into account the interactions between the genes, we consider that for all 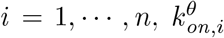 is a function which depends on the full vector *P* via the GRN, represented by a squared matrix *θ* of size *n*. We call these functions the bursts rate functions in the following. We define for all *i* = 1, …, *n*:

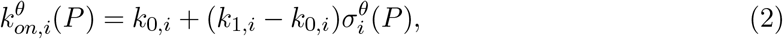

Where 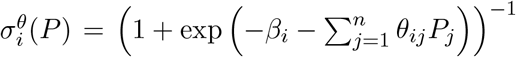. The parameter *β*_*i*_ represents the basal ac-tivity of gene *i*, and each parameter *θ*_*ij*_ encodes the interaction *j → i*. Every function 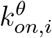 is then comprised between two positive constants *k*_0,*i*_ *< k*_1,*i*_, and 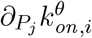 has the sign of *θ*_*ij*_. This sigmoidal form for the bursts rate functions can be interpreted as a simplification of the mechanistic form used in [16], [5].

We point out that assuming that this model is able to reproduce real single-cell datasets, we make the underlying hypothesis, which is implicit in most GRN inference methods, that, contrary to its state, the GRN structure is not modified under the action of some hidden variables. One should not that our interaction model is an approximation of the underlying biochemical cascade reactions, and that the genes we modeled need not to be transcription factors. This is possible thanks to the use of our mechanistic model which integrates the notion of timescale separation [16]. It assumes that every biochemical reaction such as metabolic changes, nuclear translocations or post-translational modifications are faster than gene expression dynamics and that they can be abstracted in the interaction between 2 genes [5]. To explicitly model slow changes like some epigenetic changes by allowing the structure of the GRN to change during differentiation, would definitely add realism but at the cost of a much higher complexity: the number of additional unobserved parameters would dramatically increase the problems of identifiability, making a reverse-engineering method out of reach.

### 1.2 Simplification in the fast transcription regime

In line with several experiments [2] [22], we consider that mRNA bursts are fast in regard to protein dynamics, *i*.*e d*_0,*i*_ » *d*_1,*i*_ with 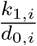 fixed. The correlation between mRNAs and proteins produced by a gene *i* is then very small, and the model can be reduced by removing mRNA and making proteins directly depend on the burst. We obtain a simplified network model in which only proteins are described:

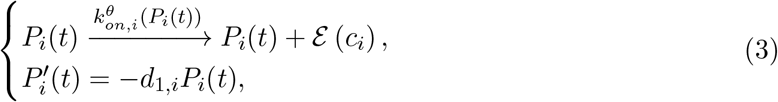

where we define 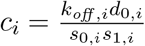.. The gene expression space Ω is then the set of possible values for the vector *P*, which is ℝ^*n*+^. The related master equation, characterizing the time-dependent distribution of *P*, turns out to be integro-differential (see Appendix B).

### 1.3 Metastability and reduction

We now perform a reduction of the model 3 into a coarse-grained model on a limited number of metastable basins, providing an approximate landscape of the differentiation process. For this sake, we introduce a typical time scale 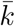 for the rates of promoters activation 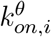, and a typical time scale 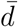 for the rates of proteins degradation. Then, we define the scaling factor 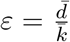 which characterizes the difference in dynamics between two processes: 1. gene bursting dynamics and 2. protein dynamics. It is generally considered that promoter switches are fast with respect to protein dynamics, *i*.*e* that *ε «* 1, at least for eukaryotes [48].

In that context, we can approximate the conditional expectation of the bursts of proteins associated to a gene *i* knowing the proteins vector *P*, that we denote *ρ*_*i*_(*P*), by its quasistationary approximation 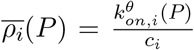. The model (3) can therefore be coarsely approximated by a system of ordinary differential equations:

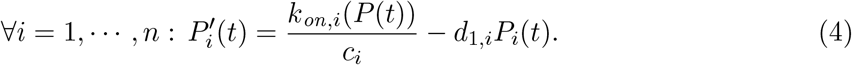

Intuitively, these trajectories correspond to the mean behaviour of a cell in the weak noise limit, i.e when bursts occur much faster than proteins concentration changes. As shown in [11] (in the general case where promoters are explicitly included in the model), for any *T < ∞*, a random path (*X*^*ε*^(*t*))_0*≤t≤T*_ converges in probability to a trajectory (*x*(*t*))_0*≤t≤T*_ solution of the system (4) when *ε →* 0.

Assuming that the dynamical system (4) has no limit cycles or more complicated orbits, the gene expression space can be then decomposed in a set of basins of attraction *Z* = *{Z*_1_, …, *Z*_*m*_*}*, respectively associated to *m* stable solutions of:

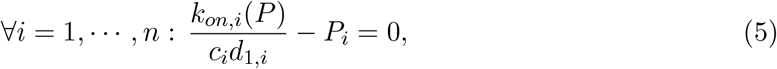

which are called the attractors of the process. Without noise, the fate of a cell is fully characterized by its initial state *x*_0_, as it converges to the attractor of the basin of attraction it belongs to, which is a single point by assumption. However, noise can modify the deterministic trajectories in at least two ways. First, in short times, a stochastic trajectory can deviate significantly from the deterministic one. In long time, stochastic dynamics can even push the trajectory out of its basin of attraction to another one, changing radically the fate of the cell in a way that cannot be catched by the deterministic limit. We illustrate in Figure 2 this situation for a toggle-switch network of two genes, where the scaling factor *ε* determines the observation of random transitions between two basins of attraction in a given time.

**Figure 2:**
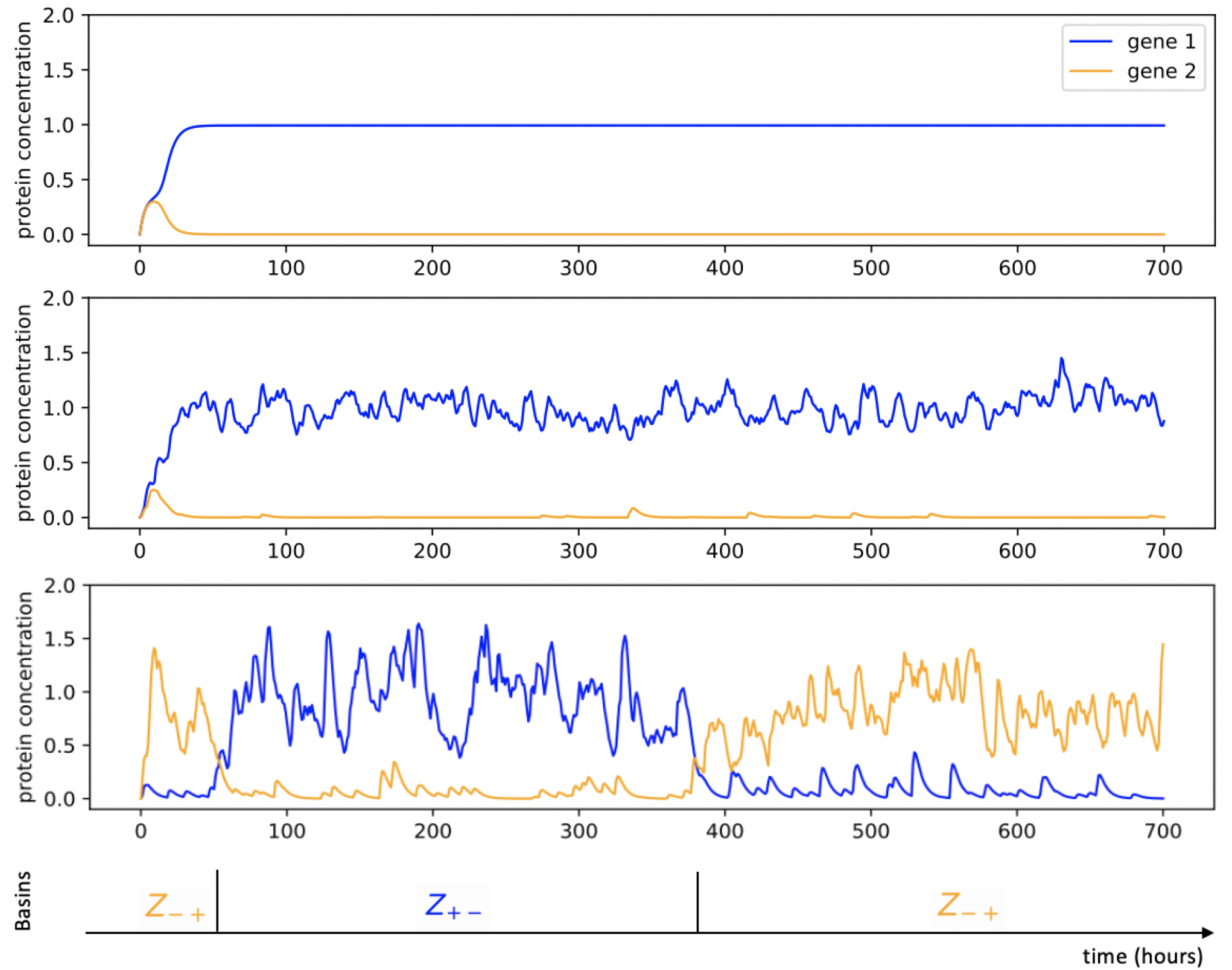
Example of trajectories associated to the symmetric toggle-switch network described in Table 1, for different values of *ε*: from top to bottom, *ε* = 0, *ε* = 1*/*50 and *ε* = 1*/*10. For *ε* = 1*/*10, we observe stochastic transitions between the two metastable basins denoted *Z*_+*−*_ and *Z*_*−*+_.

Adopting the paradigm of metastability referred in the introduction, we identify the basins of attraction associated to the equilibrium of the deterministic system (4) to cell types [18]. A cell type then corresponds to a metastable sub-region of the gene expression space, and the process can be coarsely reduced to a new (Markovian) discrete process on the cell types. These cell types represent the potential wells in the developmental landscape of differentiation associated to the model (3) [49], the centers of which are the attractors solutions of the system (5). To characterize more precisely the landscape, it would remain to describe:

1. the energetic barrier separating the cell types;
2. the curvature of the potential wells.

Point 1. corresponds to the transition rates of the coarse-grained model on the cell types, which are known to be very difficult to link analytically to a GRN [51]. They generally depends on the values of the stationary distribution *û*^*θ*^ of proteins on the saddle point of the system (4), which are located on areas of the gene expression space where the probability to find a cell is weak. As discussed in Section 6.2 with more details, this feature does not seem to be exploitable in the context of GRN inference. Point 2. can be described by the behaviour of the potential function *V* ^*θ*^ = *−* ln *uû*^*θ*^) in the neighborhood of the attractors. This feature seems more accessible because cells are likely to be measured around the attractors in the gene expression space. We are now going to develop an heuristic reasoning for approximating the function *û*^*θ*^ when the bursts rate functions are of the form (2).

## 2 From GRN to mixture approximation

### 2.1 Mixture approximation of the proteins distribution

On one side, the transitions of the coarse-grained model described in Section 1.3 happen at a time scale which is expected to be of the order of 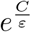, where *C* is an unknown constant depending on the basins (owing to a Large deviations principle studied in [51]). For small *ε*, such transitions are then generally rare events, and it can be considered that the process spends in each basin a time long enough to equilibrate inside, *i*.*e* that a cell reaches between each transition its quasistationary distribution within a basin. On the other side, the sigmoidal form (2) for the functions 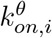 implies that these functions must not vary significantly within a basin: a rough approximation could lead to identify the bursts rate in each basin by its dominant rate inside, corresponding to the value of the function *k*_*on,i*_ on the attractor *P*_*z*_. For any gene *i* = 1, *…, n* and any basin *z ∈ Z*, we can then approximate:

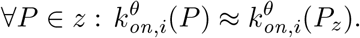

This implies that the quasistationary distribution of a cell within a basin can be approximated by the stationary distribution of the model when the burst rates are constant. The marginal distribution of each gene then appears as a Gamma distributions, which can be shown to be the unique solution of the stationary master equation of the model in one dimension when *k*_*on*_ is constant [24]. We obtain the following approximation for the quasistationary distribution 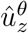 within each basin *z ∈ Z* associated to the network *θ*:

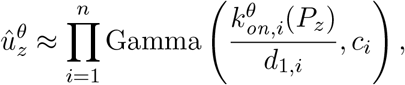

Finally, we can approximate the stationary distribution of the process associated to a given GRN *û*^*θ*^ by a mixture of Gamma distributions. Denoting for any 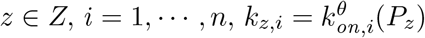, we obtain

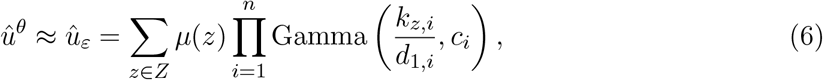

where *μ* is the probability vector on the basins at the steady-state, and we recall that *ε* depends on the values of the vectors *d*_1_ and *k*_*z*_. In some sense, this approximation consists in reducing the dependence between genes resulting from the GRN to the coexistence and relative weight of different basins corresponding to the different possible modes of promoters frequency.

The heuristic analysis presented above then states that when the functions *k*_*on,i*_ are close to constant functions inside the basins and that the scaling factor *ε* is small, the mixture approximation (6) is a good approximation of the distribution *û*^*θ*^. We remark that this is straightforward when the scaling factor *ε* is close to 0. Indeed, any GRN *θ* is associated for any value of *ε* to an unknown stationary distribution, that we denote 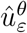, which converges to a sum of Dirac on the attractors when *ε →* 0 [36]. Then we see that the Gamma mixture *û*_*ε*_ associated to the same *ε* converges to the same sum of Dirac when *ε →* 0, as the mean of each Gamma does not depend on *ε* while the variance is proportional to *ε*: we have 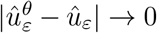 as *ε →* 0. When the noise *ε* is not negligible, the problem is more complex and finding a theoretical bound for the quantity 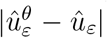, depending on *ε*, is beyond the scope of this paper. Nevertheless, we see in Figure 3 that the Wasserstein distance between the empirical distribution associated to a proteins dataset simulated from the mechanistic model (3) and the Gamma mixture distribution (6) is much smaller than the distance between the same dataset and a mixture of normal distribution (fitted with a Gaussian Mixture Model). This let us think that the Gamma mixture approximation is indeed close to the true distribution even when the weak noise limit assumption is not realistic (and that the distribution is then far from a sum of Diracs).

**Figure 3:**
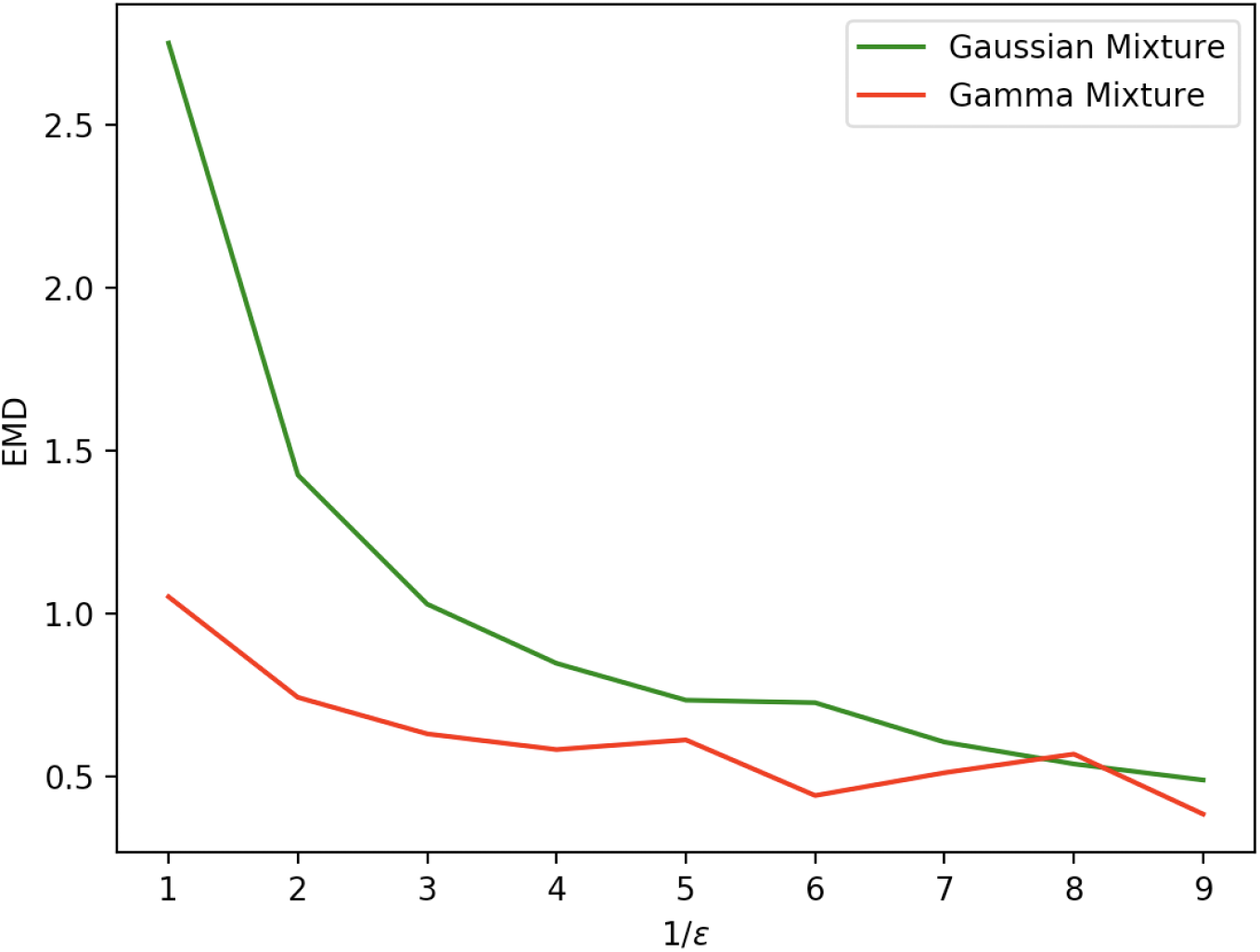
Evolution in function of 1*/ε* of the Wasserstein distances between the empirical distribution associated to a set of 2000 cells, simulated with the model (3), and both the Gamma mixture approximation (6) (in red) and a GMM approximation (in green), for the toggle-switch network described in Table 1.

### 2.2 Linking a GRN to mixture parameters

Building an analytical link between the GRN and the mixture parameters is generally out of range. Indeed, even if it was possible to solve explicitly the equation (5) defining the equilibrias associated to a GRN, it would remain challenging to study their stability in order to identify the attractors. Moreover, the probability vector *μ* is obviously linked to the transition rates of the coarse-grained model, which we recall to be very difficult to estimate from a GRN [51]. However, these parameters can be obtained with a simple numerical method, which consists in sampling a collection of random paths in the gene expression space: the distribution of their final position after a long time approximates the stationary distribution on proteins. We can then use these final positions as starting points for simulating the deterministic trajectories, given by the system (4), with an ODE solver: each of them converges to one of the stable equilibrium points. This method allows to obtain all the stable equilibria corresponding to sufficiently deep potential wells (see Figure 4(A)). Possible other potential wells can be omitted because they should correspond to basins that the process has very low probability of visiting, which do not impact significantly the coarse-grained Markov model. Repeating this operation for thousands of cells, we can find a vector *μ* describing the ratio of cells belonging to each basin (see Figures 4(B) and 4(C)). When the number and length of the simulations are large enough, the vector *μ* should be a good approximation of the stationary measure on the basins.

**Figure 4:**
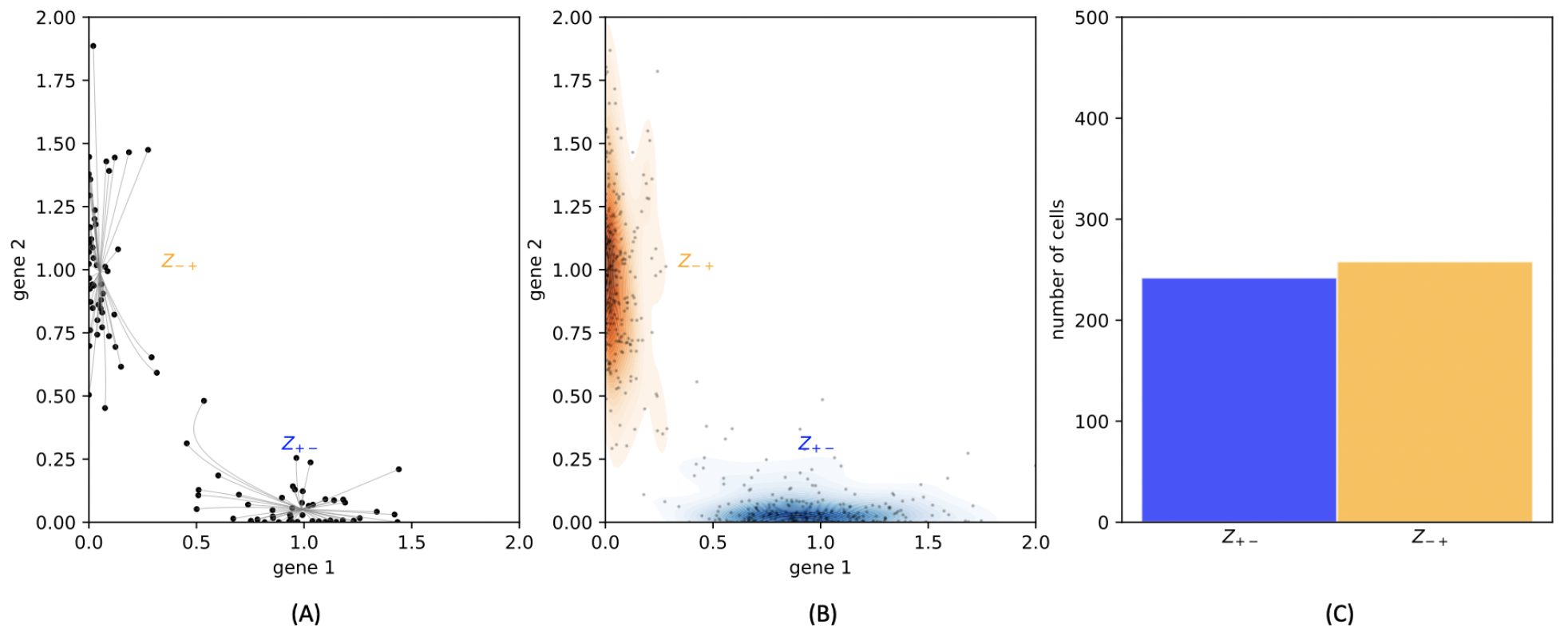
(A): 100 cells are plotted under the stationary distribution. The relaxation trajectories allow to link every cell to its associated attractor. (B): 500 cells are plotted under the stationary distribution. They are then classified depending on their attractor, and this figure sketches the kernel density estimation of proteins within each basin. (C): The ratio of cells that are found within each basin gives an estimation of the stationary distribution on the basins.

### 2.3 Mixture approximation of time-varying distribution

In the mixture approximation, the marginal on proteins of the stationary distribution of a single cell is characterized by a hidden Markov model: in each basin *z ∈ Z*, which corresponds to the hidden variable, the vector *P* is randomly chosen under the quasistationary distribution *û*_*z*_ of *X* | *Z* = *z*. Therefore, the mixture distribution can also be used as a proxy for the time-varying distributions of the bursty model (3). Denoting *µ*_*t*_ the distribution on the basins at any time *t*, and considering that a cell reaches almost immediately its quasistationary distribution after entering in a basin, we obtain the approximation:

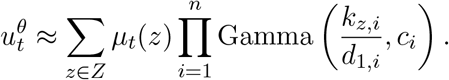

In that case, the only time-dependent parameters are the coordinates of the vector *µ*_*t*_ *∈* [0, 1]^*m*^ where *m* is the number of basins, and *µ*_*t*_(*z*) = *µ*(*z*) if *t* is such that the stationary distribution is reached.

We remark that with the method described in Figure 4, it would be possible to miss some basins that are important for describing the network, but which would not appear in the stationary distribution. It is expected to be a common situation for networks showing complex behaviours, as feedback loops or unbalanced branching structure, because the probability after a long time may be too weak for some basins to be visited, while playing an important role in the process beforehand. For such networks, the method should then be applied on a series of timepoints, and the union of all the basins identified at every timepoint should be considered (fixing the value *µ*_*t*_(*z*) = 0 for basins *z* that are not observed at time *t*). In that point of view, the basins appearing in the stationary distribution can be considered as (almost) absorbing states of the coarse-grained model.

Altogether the reduction and methods described above establish a formal basis for the definition of a simplified epigenetic landscape given a GRN, under the form of a mixture model. As the mixture parameters seem possible to obtain from a single-cell dataset (see Section 4.2), it would be fruitful to use the same formalism to assess the inverse problem of inferring the most likely GRN, given an (experimentally-determined) cell distribution in the gene expression space, a notoriously difficult task [39], [16].

## 3 Method for inferring a GRN from a mixture distribution

In this section, we discuss the problem of identifying the most-likely GRN which is associated to an approximate landscape described by the Gamma mixture distribution of the form (6). We identify such landscape to the set of parameters:

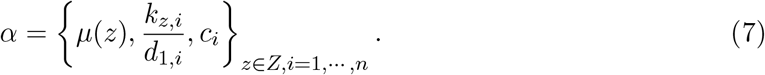

The preliminary problem of finding the most-likely vector *α* from a single-cell dataset using classical tools of statistical analysis will be developed in the presentation of the final algorithm (see Section 4.2.1).

### 3.1 Linking the GRN and the attractors

The main idea of the method consists in using the fact that all the attractors *P*_*z*_ of the bursty model verify the equation (5), *i*.*e* that for all 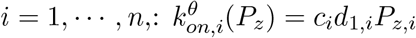. Thus, from the definition of 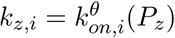, we have the relation:

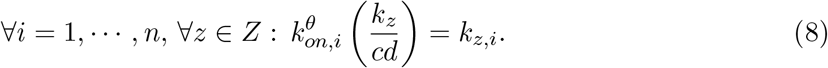

Considering that *α* is known and that we have to determine *θ*, we obtain for every gene *i* a system of *m* equations and *n* unknowns parameters corresponding to the *i*^*th*^ line of the matrix *θ*. This simple strategy may be efficient provided that there is enough basins in regards to the number of genes of interest. This is in particular the case for the toggle-switch network described in Table 1, for which its has been observed in Figure 4 that there are exactly two attractors.

**Table 1:**
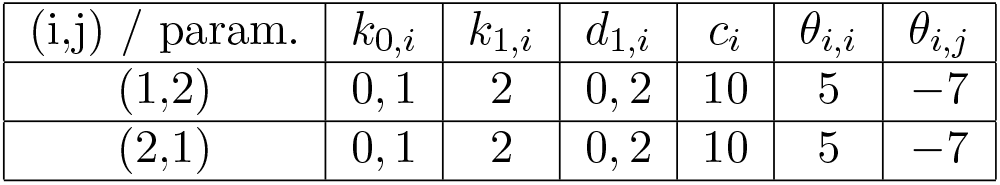
Description of the parameters of the symmetric two-dimensional toggle-switch used for the illustrations of Sections 1, 2 and Appendix C, and the network for the inference in Section 5.2 which consists in two such toggle-switch functioning in parallel.

| (i,j) / param. | $k_{0,i}$ | $k_{1,i}$ | $d_{1,i}$ | $c_i$ | $\theta_{i,i}$ | $\theta_{i,j}$ |
| --- | --- | --- | --- | --- | --- | --- |
| (1,2) | 0, 1 | 2 | 0, 2 | 10 | 5 | -7 |
| (2,1) | 0, 1 | 2 | 0, 2 | 10 | 5 | -7 |

### 3.2 GRN inference from a Gamma mixture as a minimization problem

GRN inference from a Gamma mixture is formally equivalent to identifying an adapted function *R* such that the “real” GRN *θ*^*\**^ associated to a set of mixture parameters *α* would be defined by

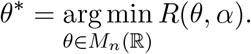

It is nevertheless important to remark that they may be no function *R* such that this equality is hold for any GRN, as a high (unknown) number of GRNs could be associated to the same mixture distribution (6). This problem is not specific to our method, and GRN inference is known to be generally a non-identifiable problem [5].

Regarding to the analysis which has been made in Section 3.1, a natural candidate for this function is

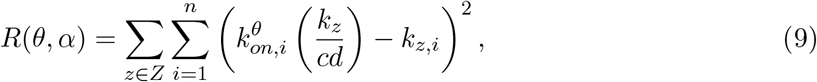

to which it may be added an adapted penalization (see Section 4.2). Taking into account the probability vector *μ* is discussed at the end of Section 4.2.2, and stems for a generalization of the method presented in this section, developed in Appendix C.

## 4 Method for inferring GRN from timestamped scRNA-seq data

In this section, we use the analysis provided in Sections 2 and 3 for developing a numerical method able to infer a GRN from scRNA-seq data, which represents the most available single-cell data at present. From a statistical point of view, scRNA-seq data gives access to the joint probability distribution of mRNAs levels associated to a given set of genes in a set of individual cells independently. Importantly, measurement techniques usually involves the physical destruction of the cell. Then, when measurements are made at several time points, for example to study convergence to a possibly new steady state after applying a perturbation to the system, we assume that the data correspond to independent samples of the marginal on mRNA of the time-varying distribution associated to the mechanistic model (1).

### 4.1 Simplified statistical model for the data

Along with the system of *n* coupled systems of the form (3) describing proteins dynamics, as presented in Section 1.2, we obtain when *ε «* 1, *i*.*e* when the bursts are frequent in regard to proteins dynamics, a quasi steady-state approximation for the conditional distribution of *M* given *P*. Under this approximation, mRNAs levels *M*_*i*_ are independent conditionally to the protein vector, and follow Gamma distributions depending on *P* :

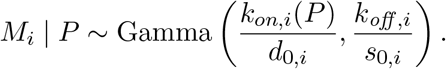

Combining this statistical model of gene expression with a Poisson model, which is claimed in [45] to adequately describe the measurement process, we then obtain the following model for a set of observed single-cell mRNA transcriptomic data:

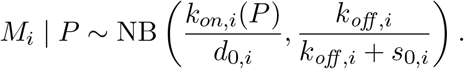

Here, NB denotes the negative binomial distribution: 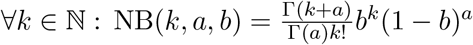.

Contrary to what has been done in [15], where the inference strategy consists in treating proteins level as latent variables and using the law on mRNAs knowing proteins as a statistical likelihood for the data, we are going to use the analysis provided in Section 2 to directly estimate the set of mixture parameters *α* from the data without the need of proteins quantity. The main idea is the following. On one side, the only information on the proteins that can be obtained is contained in the value of *k*_*on,i*_(*P*). On the other side, proteins being known to be less noisy than mRNAs, a vector *P* characterizing a cell in the gene expression space is going to be generally close to one of the attractor. Moreover, the functions *k*_*on,i*_(*P*) are almost constant within each basin, and in particular around their attractor. Thus, the precise knowledge of the proteins quantity can be considered out of reach from scRNA-seq data, and the best information on proteins we can get from the marginal distribution on mRNAs of a gene *i* is the set of the modes of bursts frequency normalized by the degradation rate, that is 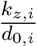.

We finally derive a simplified statistical model, making appear the distribution of mRNA counts as a mixture of negative binomial:

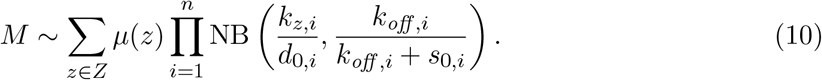

### 4.2 Inference procedure

We now describe the inference procedure for a set of *l* timestamped scRNA-seq data 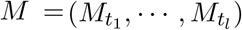, where each 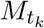 is a matrix of gene expression containing the 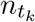 cells 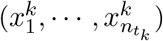. We decompose the algorithm in 2 steps:

1. A clustering step, for identifying the set of parameters 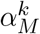 associated to the data at each timepoint *t*_*k*_;
2. A regression step, consisting in identifying the most-likely GRN *θ* associated to the sets 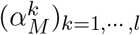, by successively solving a set of regression problems.

#### 4.2.1 Clustering

From the statistical model (10), the first step consists in inferring for each timepoint *t*_*k*_ the set of parameters

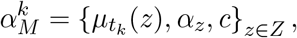

where 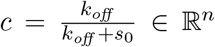 and for every 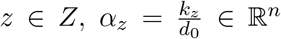. The number of genes being potentially large, a minimization of the maximum likelihood function *f* on the gene expression matrix 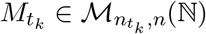, defined by

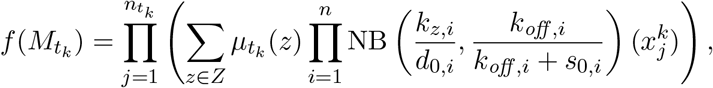

even using an EM algorithm or an other variational method, would be very uncertain, such algorithms being known to be trapped into local minima [10]. The number of components of the mixture, *i*.*e* the size of the set *Z*, remains also unknown. A good method for fitting 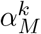 would be to apply a Bayesian procedure like a Reversible-Jump Monte-Carlo Markov Chain (RJMCMC) algorithm [42] to the whole dataset. We propose a slightly different approach, which overcomes the numerical limits of such Bayesian method on the multivariate procedure, consisting in two steps:

1. We first identify the modes of the bursts frequency which are associated to every gene *i*, independently, for the whole set of cells *M*. For this sake, we use a RJMCMC algorithm which is inspired from [6]. We obtain the values of 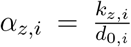. Each cell 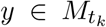, represented by a vector in ℕ ^*n*^, can be then associated to a discretized representation of the form: 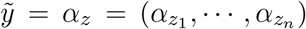, where every *z*_*i*_ characterizes one of the modes associated to the gene *i*;

2 For each timepoint *t*_*k*_, we group together the vectors 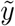 which are equals. The different groups that are obtained correspond to the modes (*α*_*z*_)_*z∈Z*_ observable at *t*_*k*_. The relative weights associated to these modes provide an estimation of the probability vector 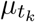.

This simplified method, in which the coupling between the different genes is taken into account only in a second stage, is likely to provide more basins than the expected number. However, we will see in Section 4.2.2 that this may be corrected by a simple modification of the cost function used in the regression step. We also discuss in Section 5.5 the fact that the number of modes associated to each gene is generally limited to 2 in the case of the sigmoidal bursts rate functions (2): then each cell *y ∈ ℕ*^*n*^ is generally projected in step 1. in a reduced discrete space of dimension 2^*n*^.

#### 4.2.2 Regression

First, in order to reduce the number of parameters and since protein levels *P*_*i*_ are not observed, we can arbitrarily set:

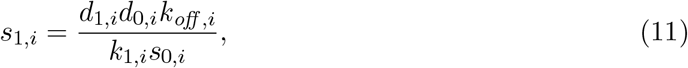

which leads to 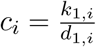. Injecting this value in the formula (5), the attractors are then defined for any *z ∈ Z* by the formula: 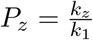. It is worth noticing that the choice of this scaling may not affect the accuracy of the inference. Indeed, the value of the parameter *s*_1,*i*_ only affects the number of proteins *P*_*i*_ that are created when *M*_*i*_ is positive: the value of *P*_*i*_ is then proportional to *s*_1,*i*_. As every parameter *θ*_*ij*_ affects the dynamics of the process through the value of the product *θ*_*ij*_ *× P*_*i*_ in the function *k*_*on,j*_, considering such scaling for *s*_1,*i*_ will only affect the scaling for the interaction coefficients *θ*_*ij*_, but not the GRN topology.

The clustering step only allows to find the values of the vectors 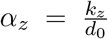 rather than *k*_*z*_. This is not a problem since the fraction which characterizes *P*_*z*_ allows to simplify the term *d*_0_, provided that 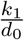 can be computed. We therefore consider that each gene reaches its minima and maximal frequency through the timepoints, at least for some cells. Thus, we define for every gene 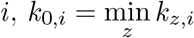 and 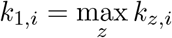, and equivalently:

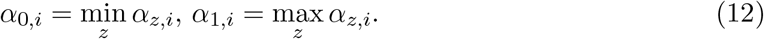

We finally obtain a characterization of the attractors which is directly accessible from scRNA-seq data:

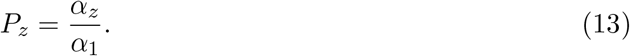

Injecting (13) in the definition of *R* given by (9), we obtain the following characterization of a cost between a GRN *θ* and a vector *α* describing the approximate landscape of the model (1):

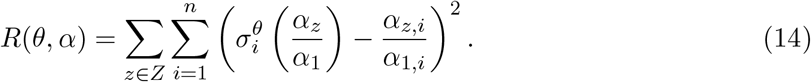

In order to define a regression problem on the variable *θ* which may associate a vector *α* obtained in the clustering step (see Section 4.2.1) to a GRN, we add three modifications to the cost function *R* given in (14).

First, we solve the problem of the possibly too high number of basins due to the clustering method, that we mentioned in Section 4.2.1, by taking into account the weight of the basins in the cost function. Then, given that we have enough cells that are detected in existing basins, the “false” basins detected in the clustering step, which should not be associated to many cells, would almost not affect the inference.

Second, in line with the idea that missing interactions is preferable to inferring false interactions between genes, we decide to use a LASSO penalization, which is known to enforce the sparsity of the network. We also add a custom penalization to deal with oriented interactions. Indeed, for every pair of nodes *{i, j}* there are two possible interactions with respective parameters *θ*_*ij*_ and *θ*_*ji*_, but it is likely that only one is actually present in the true network. Our method is likely to favor symmetric interactions because when an interaction is present in the network (e.g *θ*_*ij*_ *>* 0), then gene *j* is generally upregulated in the same time than gene *i* and it is hard to distinguish whether *θ*_*ji*_ *>* 0 or not. Then we want these two interaction parameters to compete each other, such that only one is nonzero after the regression step, unless there is enough evidence in the data that both interactions are present. We obtain the following penalization

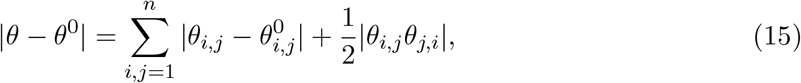

where *θ*^0^ is defined as the null matrix of size *n* if there is no prior information on the GRN. The coefficient in front of the product | *θ*_*ji*_*θ*_*ij*_ | is chosen small enough to ensure that if both *θ*_*ji*_ and *θ*_*ij*_ have been detected at a timepoint, it would generally cost more to put one back close to 0 than to keep it at its computed value.

Third, the temporal dynamics of the process, when available, must be considered. As developed in Section 2.3, the basins that have to be taken into account are all the basins identified during the differentiation process, *i*.*e* in the different snapshots of the time-varying distribution. But three principal reasons lead to consider that we should not try to infer the network from all the basins at the same time:

1. We can generally see the effect of a GRN in a cell as a signal which is transmitted to the genes [5]. Before the signal has been completely transmitted in the network, many genes are likely to be in a state which does not reflect the GRN. At these moments, the cell is far from the equilibrium. For this reason we would like to take into account the first timepoints, where some genes have not seen the signal, in a weaker way than the last timepoints, where the signal has been well transmitted to all the genes.
2. A population of cells is likely to be more often observed far from the attractors before it has reached its equilibrium than after reaching it. This is due to the fact that when the distribution is far from the equilibrium, some shallow or even unstable basins can be explored, from which the cell can easily escape. In particular, the regions around bifurcations in the cell lineage are expected to be often identified as shallow basins. These basins should then almost be erased if deeper basins are explored afterwards. This also suggests that the first timepoints have to be taken into account in a weaker way than the final ones.
3. Finally, recalling that GRN inference is known to be generally a non-identifiable problem, we simply cannot neglect the information that is brought by temporal information when it is available.

We thereby adopt an iterative approach: for each timepoint, the network is actualized by minimizing the function (14) with the penalization (15), taking as initial condition the network inferred at the previous timepoint. Then, we do not penalize the network itself but its variations to the initial condition. This should satisfy the points 1. and 2., because an erroneous interaction that would have been catched at an early timepoint would be conserved in the final network only if it does not appear in contradiction with the interactions that are inferred afterwards.

Finally, the regression step consists in solving successively, for *k* = 1, *…, l*, the problem

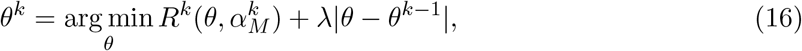

where *θ*^0^ is defined as the null matrix of size *n*, |*θ − θ*^*k−*1^| is given by (15), *λ* is a penalization coefficient and the function *R* is defined for every *θ* and 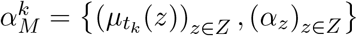 by:

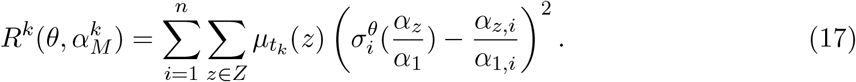

The procedure is illustrated in Figure 5. For the applications presented in Section 5, the value of the coefficient *λ* has been calibrated in order to be optimal for various datasets simulated from randomly-generated tree-like networks, like those studied in Section 5.6.

**Figure 5:**
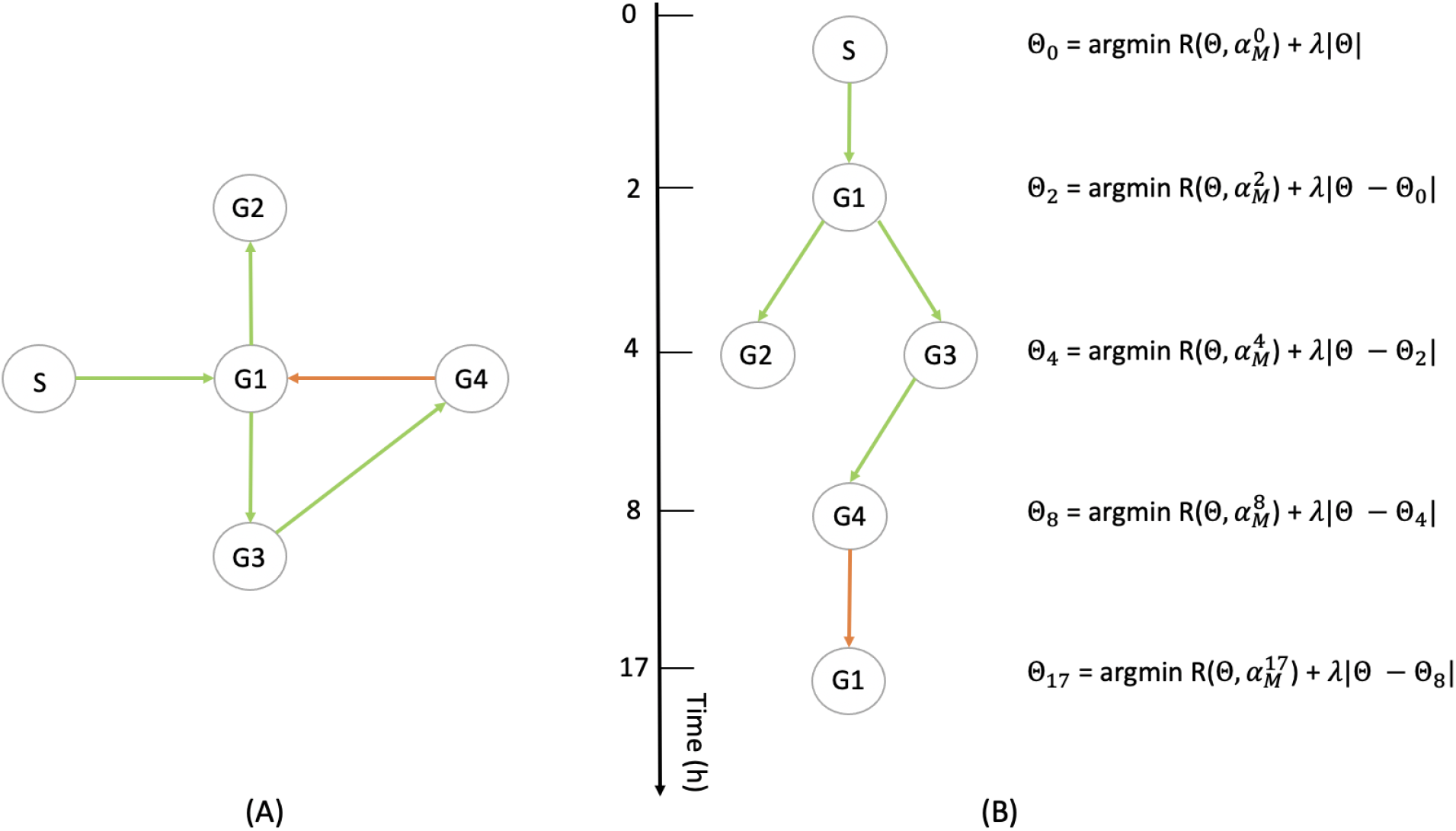
(A) Example of a 4-genes network (G1 to G4) with a stimulus (S) (see Section 5.1). (B) Illustration of the method described in Section 4.2.2: the interactions being observable only at some particular timepoints, they are progressively inferred, each optimization taking into accounts the interactions that are observed on the previous ones.

Interestingly, the cost function (17) can be obtained as the particular case of a more general method for linking a GRN to a set of mixture parameters in the case where single-cell proteomic data were available, and that mRNAs are seen as a proxy for the proteins level. We present this method in Appendix C.

We underline the fact that the function *σ*^*θ*^, defined in (2), depends only on the GRN *θ* and 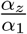, which can be directly estimated from scRNA-seq data, using the method described in Section 4.2.1 and the relations (12). Thus, we do not need to make any assumption on the value of the hyperparameters of the model, not even the ratio 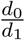 as it was the case in [15]. This is due to the fact that we do not infer the protein distribution but only the values of its main modes, which are completely characterized by the mean of the proteins distribution within each basin, and do not depend on its variance.

The two-steps method presented in Sections 4.2.1 and 4.2.2, that we call CARDAMOM (Cell types Analysis from scRna-seq Data Achieved from a Mixture MOdel), shows good results for the simulated datasets that are presented in Section 5.

#### 4.2.3 Back to the model: consistency of the algorithm and verifications

The inference method presented above corresponds to the calibration of the mechanistic model (1) driven by burst rates functions of the form (2). Recalling that GRN inference is generally not an identifiable problem, many networks are likely to reproduce a dataset. Beyond the topology of the network that we expect to be well reconstructed by the algorithm, a GRN given by CARDAMOM should be considered as accurate if it allows to reproduce the datasets used for the inference, seen as partial observations of the time-varying distribution associated to the mechanistic model. It would then be natural to use the quantitative parameters inferred with CARDAMOM to simulate new snapshot data that we could compare to the original data. However, we point out the fact that the inference method described in Section 4 only used the attractors of the metastable basins associated to the deterministic limit (4), while we recall that the dynamics of the model is mainly determined by the transition rates of the coarse-grained model on the basins, which depends on the value of the potential on the saddle points of the system (4) rather than the attractors. This is in line with the fact that the vector *α* is supposed to approximate accurately the potential wells of the landscape but not the energetic barrier between them, which remains out of reach. Thus, the method is supposed to find a network which should allow to recover the same metastable basins than the ones observed in the data, the order in which they are visited, but not the right temporal dynamics.

In order to estimate the quality of the inference while taking into account the limitations mentioned above, we should simulate the model (1) for comparing both the initial and the long-time distribution of the model associated to the inferred GRN with the first and the latest snapshot which is available on the data, and verify that simulated cells have a transitory evolution similar to the data (in terms of distribution), even if not necessary with the same temporality. This is the object of Section 5.4.

## 5 Results

In this section, we present the performance of CARDAMOM on simulated datasets from 3 different types of networks. For each network, the datasets are obtained by sampling independent cells at a certain number of timepoints after applying a stimulus.

### 5.1 Simulating data with stimulus

The simulations of the model (1) are performed with the python package HARISSA presented in [15], which is based on an efficient thinning method for simulating the model (1). For reproducing *in vitro* differentiation processes, we first simulate every network until reaching a first equilibrium before *t* = 0. At *t* = 0, we introduce a new gene called “stimulus”, the proteins level of which is artificially maintained at the value of 1. This value corresponds to to its highest possible attractor: indeed, the proteins scaling (11) implies that no coordinates of the attractors can be greater than 1 in the gene expression space. Proteins being much less noisy than mRNA (i.e 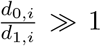), the protein level associated to a gene is expected to be close to the attractor associated to the basin it belongs to, and forcing a gene to be fully expressed in the model (1) with the scaling (11) is equivalent to force its value to 1. Then, the stimulus represents the effect of an artificial perturbation in the differentiation environment inducing cells to evolve towards a new equilibrium. The datasets correspond to the sample of independent cells at a sequence of timepoints.

### 5.2 Inference of a toggle-switch network from a static expression data sampled under the long time distribution

We first evaluated the ability of CARDAMOM to find the parameters of a 4-genes network consisting in 2 independent toggle-switch of the same form than the one described in Table 1. To this aim, we simulate 10 dataset by sampling 50 independent cells at 2 timepoints 0 and 20*h*. In that case, the stimulus has no effect on the system. Inference is then independently performed for every datasets and the mean of all the significant values obtained for each network is presented in Figure 6, compared to the network used for the simulations. We observe in Figure 6 that the algorithm allows to reconstruct very well the topology of the network, detecting which pairs of genes are in relation together and the sign of the interactions. All other coefficients of the GRN matrix are smaller than 0.3, which can be considered as a residual noise with regards to the principal edges. However, the algorithm computes the values of the coefficients up to a multiplicative factor, which is in line with the limitations we pointed out in Section 4.2.3. We remark that the algorithm slightly overestimates the diagonal coefficients, which correspond to self-regulation parameters of the genes : the problem of the inference of these parameters being notoriously difficult to infer reliably [40], we will not take into account the diagonal coefficient of the GRN matrix for evaluating the algorithm performances in the following.

**Figure 6:**
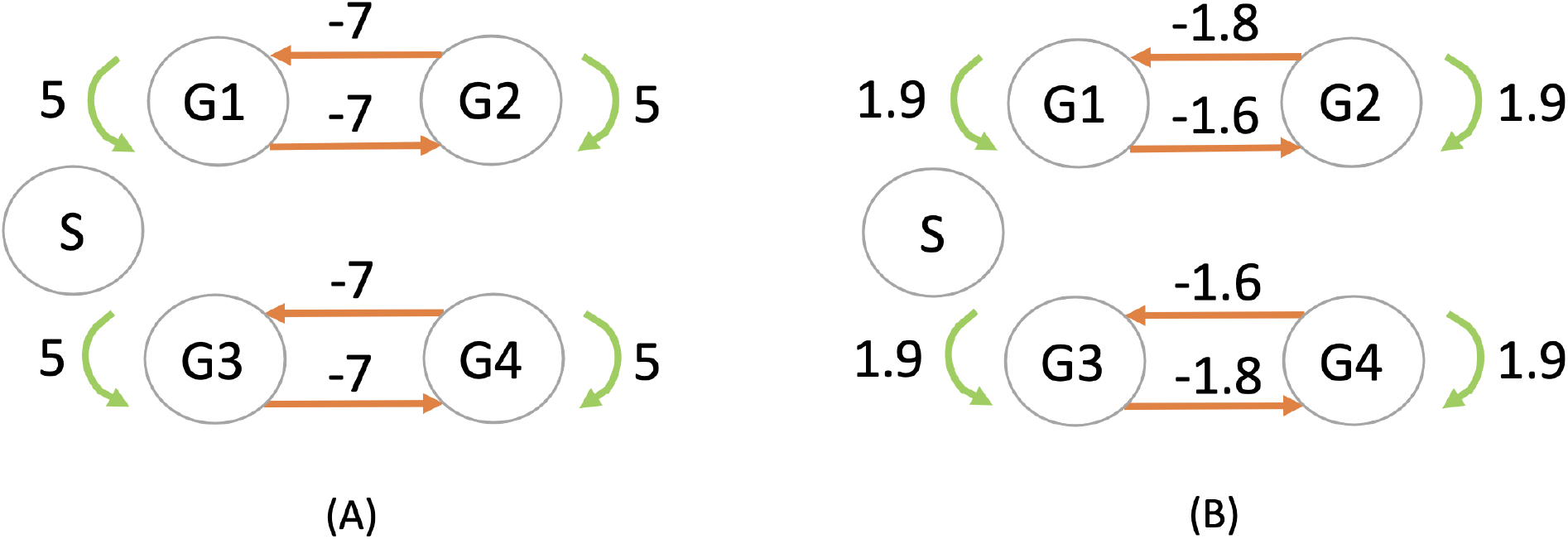
Inference of a 4-genes network (G1 to G4) consisting in two independent symmetric toggle-switch networks with parameters described in Table 1. The stimulus (S) has no effect on the network, but is nevertheless represented in order to verify whether the inferred network takes it into account or not. The network used for simulating the datasets in (A) is compared with the network inferred by CARDAMOM in (B).

### 5.3 Inference of a 4-genes network with branching and feedback loop from timestamped data

We now evaluate CARDAMOM on the 4-gene network described in Figure 5 and Table 2. Although such a small network may appear very simple, it already has some interesting features (branching, feedback loop with inhibition) and is interesting for inference. To this aim, we simulate 10 datasets by sampling independent cells at 10 time points *t* = 0, 2, 4, 6, 8, 11, 13, 15, 17, and 20*h*, with 50 cells per timepoint (then 500 cells in total for each dataset). Inference is independently performed for every dataset and the results are merged into receiver operating characteristic (ROC) and precision-recall (PR) curves that are shown in Figure 7. CARDAMOM turns out to reconstruct very efficiently the topology of the network. We precise that the sign of the interactions that are inferred is well preserved comparing to the original network.

**Table 2:** Description of the parameters of the 4-genes network used for the inference in Section 5.3. All others parameters are similar than for the toggle-switch network of Table 1, the same for every genes. The *i*^*th*^ line (resp. the *i*^*th*^ column) of the matrix *θ*, correspond to the influence of every genes on the gene *i* (resp. the influence of the gene *i* on every genes).

| $\theta_{ij}$ | $\theta_{.,0}$ | $\theta_{.,1}$ | $\theta_{.,2}$ | $\theta_{.,3}$ | $\theta_{.,4}$ |
| --- | --- | --- | --- | --- | --- |
| $\theta_{0,.}$ | 0 | 0 | 0 | 0 | 0 |
| $\theta_{1,.}$ | 10 | 0 | 0 | 0 | -10 |
| $\theta_{2,.}$ | 0 | 10 | 10 | 0 | 0 |
| $\theta_{3,.}$ | 0 | 10 | 0 | 10 | 0 |
| $\theta_{4,.}$ | 0 | 0 | 0 | 10 | 0 |

**Figure 7:**
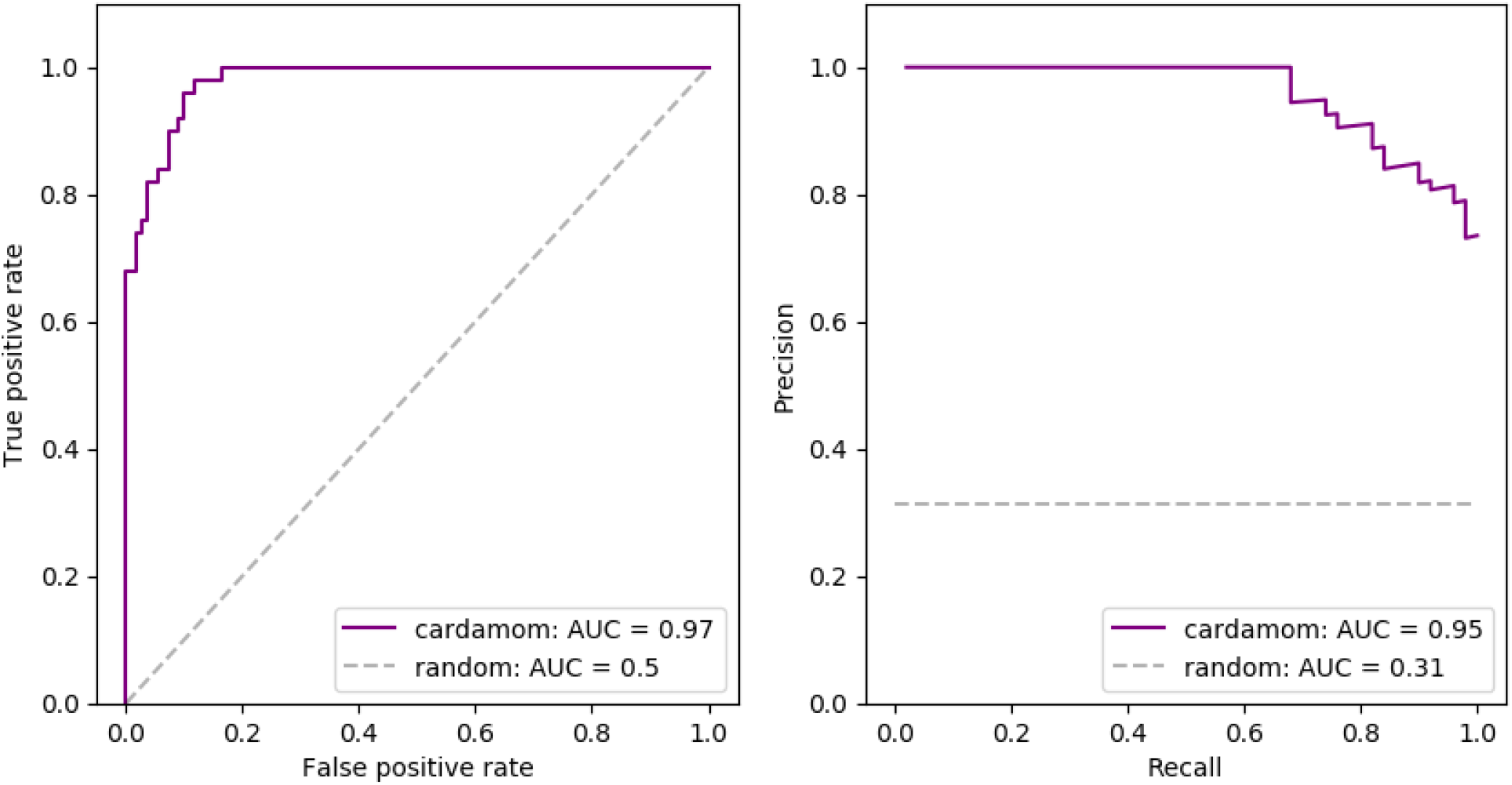
Inference results for the 4-gene network described in Table 2. Performances are measured in terms of receiver operating characteristic curves (ROC) and precision-recall curves (PR) obtained for 10 independently simulated datasets. Each dataset contained the same 10 timepoints and 50 cells per time point. The dashed gray line indicates the average score that would be obtained by the random estimator (detecting a link or not with equal probability).

**Figure 8:**
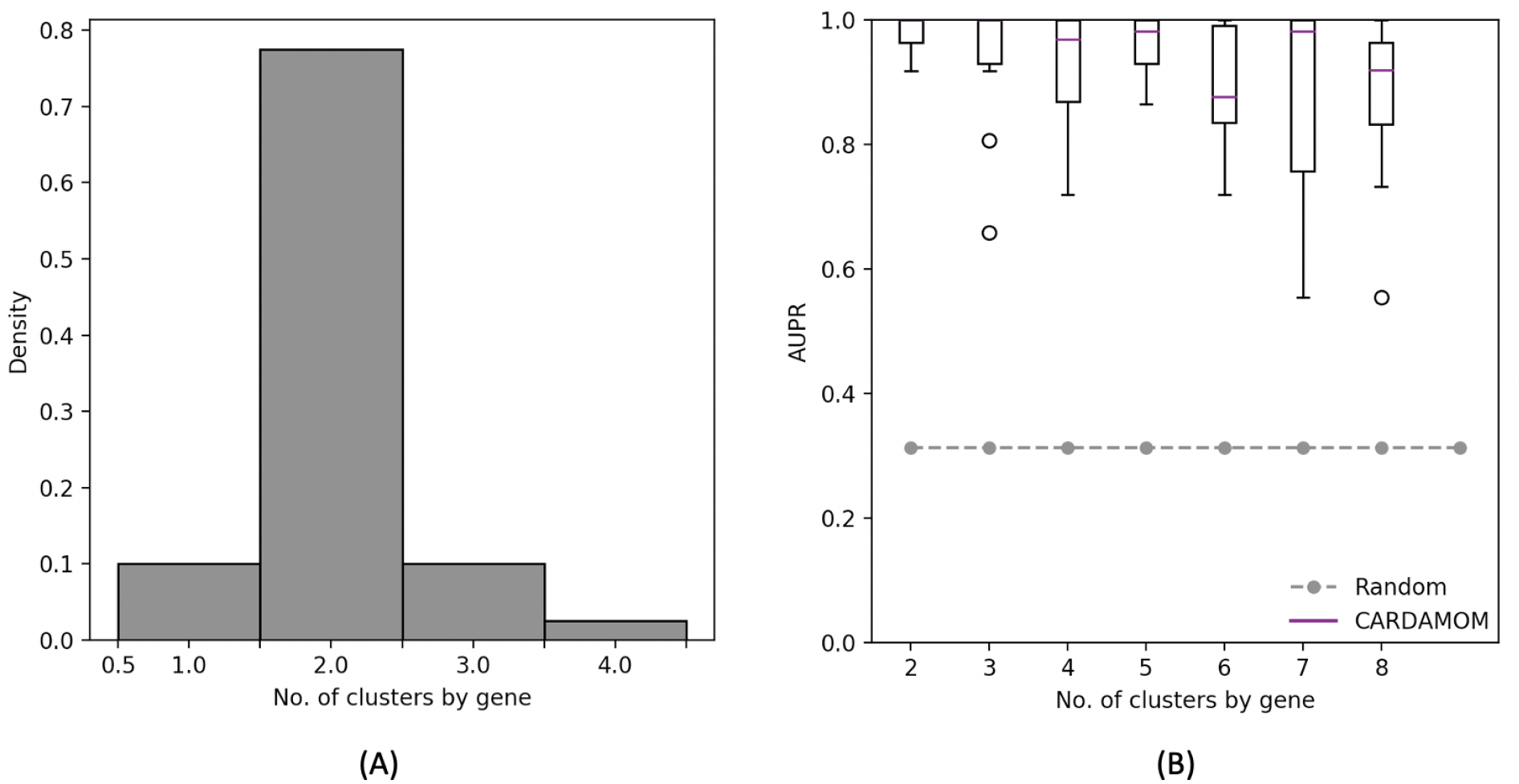
(A): Distribution of the number of clusters found by the RJMCMC algorithm during the clustering step for each gene of the 4-genes network described in Table 2. (B): Comparison of the performances of CARDAMOM for the same network, when imposing different values for the number of clusters in the clustering step. Performances are measured in terms of area under precision-recall curve (AUPR), based on 10 datasets corresponding to the same network.

### 5.4 Reproduction of the data

The bursty model (1) with *k*_*on*_ functions defined by (2) can satisfactorily produce expression data for which marginal distributions of genes are close to mixtures of negative binomial distributions, which is known to be biologically relevant. In line with what was discussed in Section 4.2.3, we do not expect the temporal dynamics associated to the network inferred by CARDAMOM to be synchronised with the original data. We then focus on the comparison between the initial and final distributions of the model when simulated by the network presented in Table 2 and the network inferred in Section 5.3, observed at respectively 0*h* and 20*h* after applying the stimulus (see Figure 9). We underline the fact that we do not take the initial distribution of the data to initialize the simulation, but we rather sample a distribution using the inferred network with the stimulus fixed to 0.

**Figure 9:**
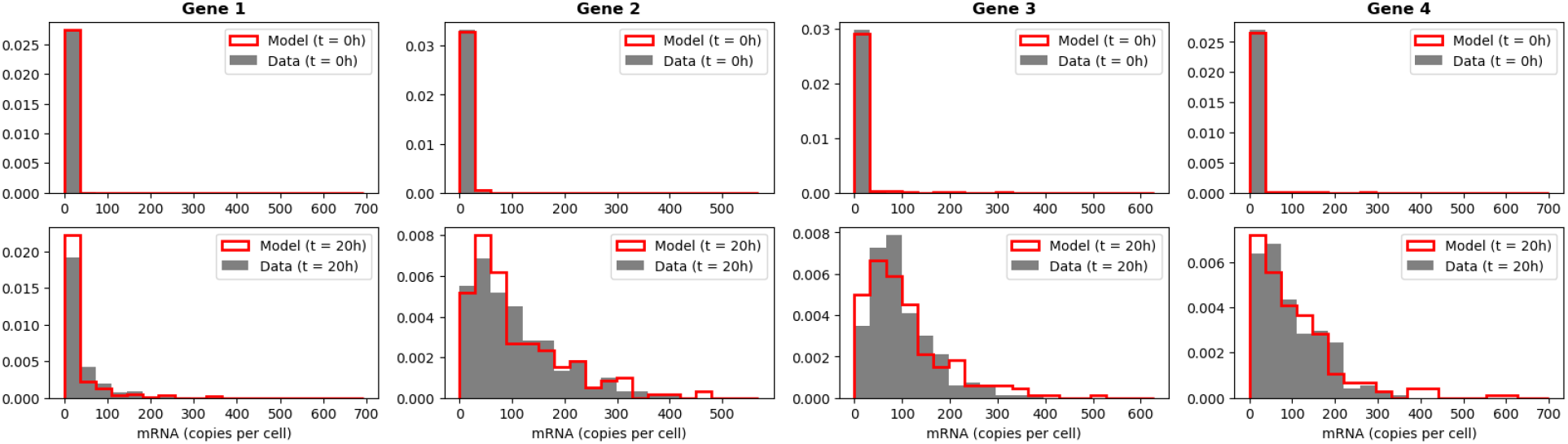
Comparison of the initial (*t* = 0*h*) and final (*t* = 20*h*) empirical marginal distributions of 4 genes simulated by the mechanistic model with the GRN described in Table 2, and the dataset simulated from the GRN inferred by CARDAMOM from the previous dataset (with 200 cells per timepoints).

We also compare in Figure 10 the two temporal dynamics of the two complete datasets, one used for the inference and one simulated with the inferred network. The data are projected altogether with the method *umap* [27] and we show on each subfigure the cells corresponding to one of the datasets. These cells are then classified depending on the time where the measures are realized after applying the stimulus. The results suggest that the model has been relatively well calibrated through the inference procedure.

**Figure 10:**
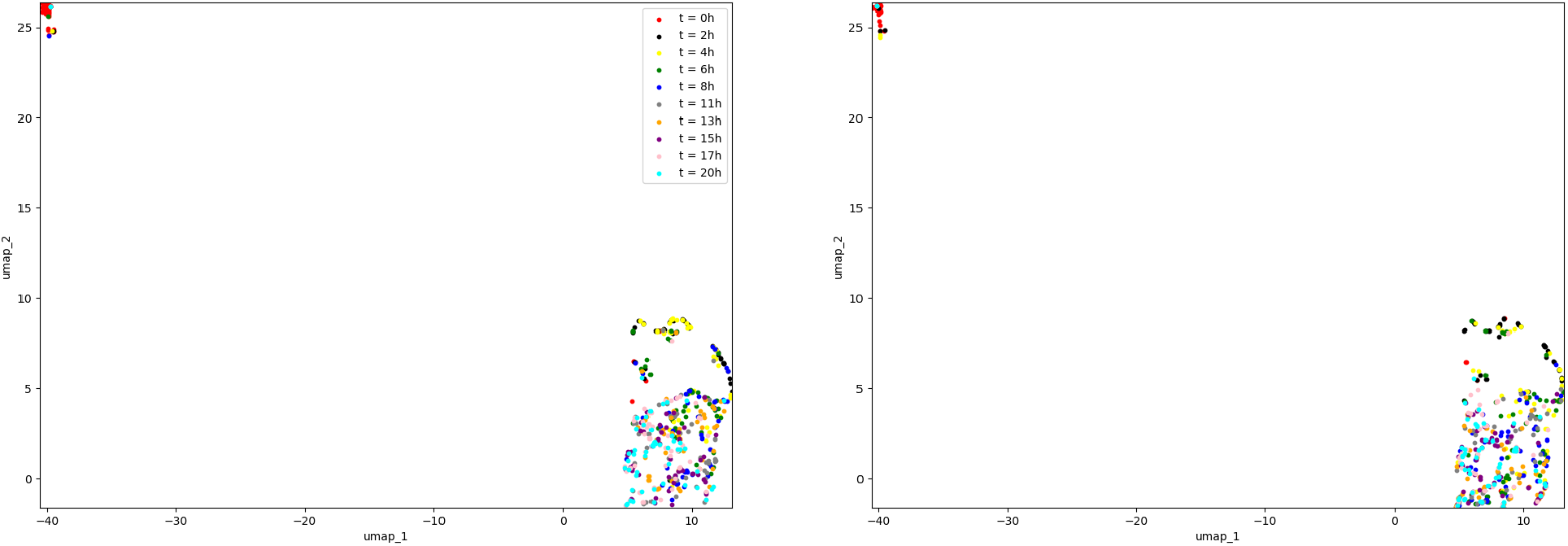
Representation in 2 dimensions of (A) the dataset used for the inference, simulated from the GRN described in Table 2 and (B) the dataset simulated from the GRN inferred by CARDAMOM. Each dataset contains 500 cells divided in 10 timepoints.

### 5.5 Simplification of the method for sigmoidal burst rate functions

The clustering method described in Section 4.2.1 allows to take into account any number of modes for the burst frequency associated to each gene. However, it is natural to ask if such complexity is compatible with the one allowed by the choice of the functions *k*_*on,i*_. In fact, when these functions have the sigmoidal form described in (2), they cannot be expected to be associated to a high number of attractors with intermediate values comprised between *k*_0,*i*_ and *k*_1,*i*_. A rough simplification consists in considering that every gene is the result of a mixture of the form (6) with only two modes, which can be expected to be close to the values of *k*_0,*i*_ and *k*_1,*i*_. Following this idea, the clustering step of CARDAMOM can be replaced by a simple binarization of the cells, where the mRNA value measured for each gene is classified as belonging either to the mode *k*_0,*i*_ or *k*_1,*i*_, with a likelihood ratio between the two negative binomials associated to these parameters. We underline that this approximation does not mean that the data always exhibits strong bimodality. It only means that the fact that they are sometimes observed far from their extremum, which is expected at least during the period of transition after applying the stimulus, does not mean that these transitory states are stable.

The simplification consisting in imposing a number of clusters equal to 2 for each gene would not affect the efficiency of the algorithm for the 4-genes network studied in Section 5.3. Indeed, we observe in Figure 8(A) that the RJMCMC algorithm applied to each gene of the network for every datasets simulated from this network usually computes a number of clusters equal to 2, and in Figure 8(B) that imposing a number of cluster greater than 2 makes slightly decreases the efficiency of the algorithm. We discuss in Section 6.4 the consequences of the accuracy of such simplified method, which may argue in favor of more complex burst functions than the ones of the form (2), or even adaptive functions depending on the clustering step described in Section 4.2.1.

### 5.6 Application to larger networks

We now consider the cases of tree-like activation networks of 5, 10, 20, 50 and 100 genes. For each case, we simulate ten datasets corresponding to 10 random networks of the same size, that sampled from the uniform distribution over trees rooted in the stimulus [15]. All datasets contain the same 10 timepoints *t* = 0, 2, 5, 8, 11, 13, 16, 19, 22, and 25*h*, with 100 cells per time-point (then 1000 cells in total for each dataset). Inference is then independently performed for all dataset with CARDAMOM and the state-of-the-art method GENIE3 [19]. The results are presented in terms of Area Under Precision-Recall curves (AUPR), in Figure 11A). CAR-DAMOM turns out to reconstruct more efficiently the topology of the network than GENIE3. The strong decrease in performance when the number of genes gets higher is due not only to the lack of data per timepoints in regards to the number of interactions, but also to the choice of the timepoints. Indeed, a sequence of timepoints that is too coarse to catch the dynamics would lead to a lack of accuracy in the inference, and a sequence which is too tight would often be too short, leading to miss the activity of some genes. This appears to be the case from 20 genes in Figure 11A).

**Figure 11:**
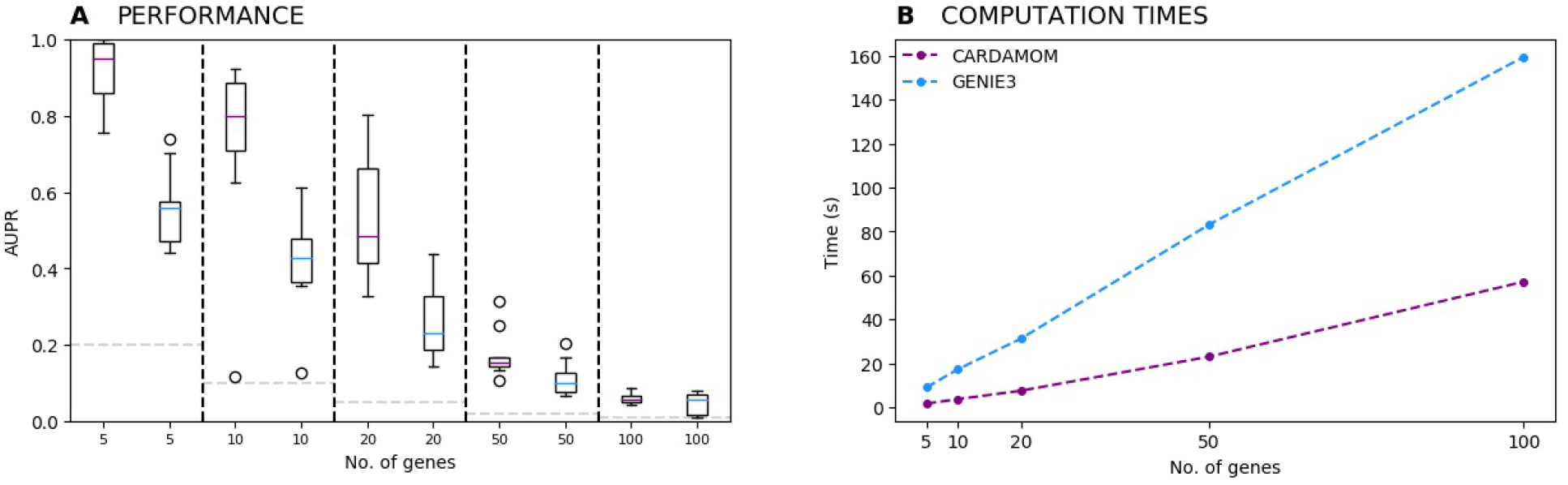
(A) Evolution of the performances of CARDAMOM and GENIE3 when increasing the number of genes. For each number of genes, we represent a boxplot of the AUPR scores computed for ten datasets simulated with 10 different randomly generated tree-like networks. (B) Evolution of the average computational time measured for inferring the tree-like networks with respect to the number of genes.

For every size of network, an average runtime is obtained after inferring the 10 datasets associated to the 10 tree-like networks, on a 16-GB RAM, 2,4 GHz Intel Core i5 computer. We see in Figure 11B) that for both algorithms the computational speed increases linearly with respect to the number of genes, with a slope which is significantly lower for CARDAMOM. These results show that the algorithm is suitable for realistic number of genes and cells.

## 6 Discussion and prospects

In this discussion, after clarifying in Section 6.1 the importance of the clustering step of CAR-DAMOM, we go deeper in Section 6.2 into the link between the mixture approximation and the popular notion of Waddington landscape. Then, we discuss in Section 6.3 the applicability of our method to real single-cell datasets. Finally, we show in Section 6.4 that the regression step of the method highlight the link existing between the mechanistic model and a neural network, and in what way this analogy paves the way for a more general method using adaptive bursts rates functions.

### 6.1 Importance of the clustering

The attentive reader should have remarked that the function (17) which is optimized in the second step of CARDAMOM appears very simple and close to a generalized linear model, which is surprising for an analysis based on a mechanistic model. It is then natural to ask whether the clustering step remains important or if it is possible to reduce the method to the regression step by identifying each cell to a distinct attractor without losing an important accuracy (the clustering step would then simply consists in rescaling each count of a gene *i* by the parameters *c*_*i*_). We compare in Figure 12 this coarse simplification with CARDAMOM on the datasets used in Section 5.3. We observe that such simplification generates a significant decrease in accuracy, confirming that the method cannot be reduced to such generalized regression.

**Figure 12:**
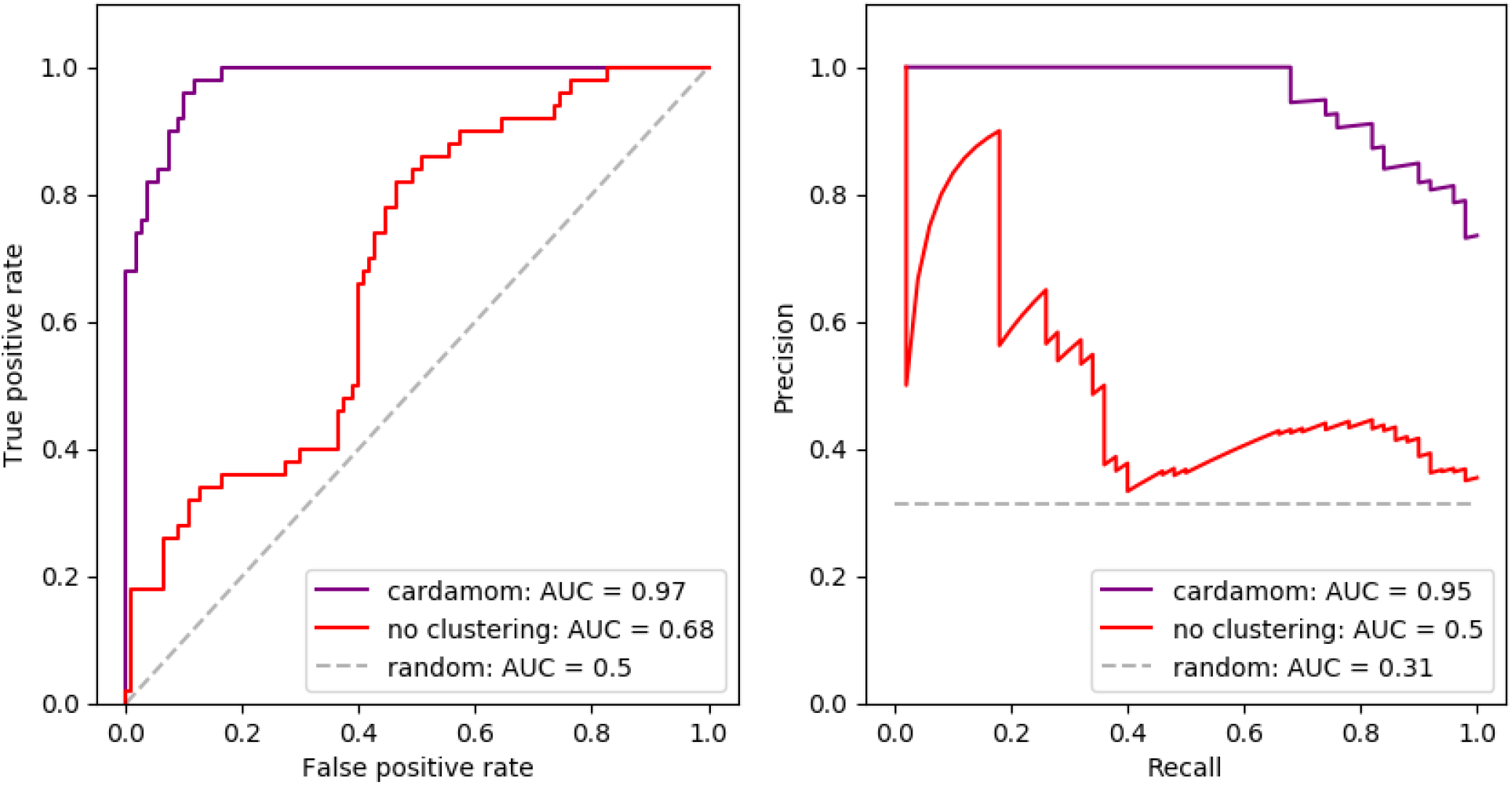
Analogy of Figure 7, where we compare (A) the ROC curves and (B) the PR curves of the GRN reconstruction performed by CARDAMOM with clustering (in purple) and without clustering (in red, considering that there are as many clusters as cells in each dataset).

### 6.2 Limits of the method for finding the developmental landscape

In this section, we come back on the Gamma mixture approximation (6) and its limitations for describing the developmental landscape of differentiation, in order to remove some of the confusion that might arise from misuse of this approximation. As we presented in Section 2, this mixture approximation is based on two assumptions:

1. The functions *k*_*on,i*_ are close to constant inside the basins:
2. The jump of a cells from one basin of attraction of the deterministic limit (4) to another is a rare event.

Then, in terms of Waddington landscape, the mixture is supposed to describe accurately the bottom of the potential wells associated to a GRN and, at least under the assumption 1., the curvature of the potential wells that are deep enough. However, it does not describe accurately the boundaries of these wells, which characterize the energetic barrier and correspond precisely to areas of the gene expression space where the functions *k*_*on,i*_ have an important gradient, making the approximation not accurate. In other words, the potential

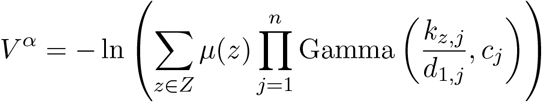

is not supposed to be a good approximation of the real potential *V* ^*θ*^ = *−* ln (*û*^*θ*^) associated to a GRN when a cell is far from the attractors. This explains, with a statistical physic point of view, why our algorithm is not able to catch the temporal dynamics of the data, but only the same sequence of temporal distributions.

This limitation is very hard to overcome. For this sake, we should either get knowledge on the values of the cells distribution on saddle points of the landscape which are located in areas of the gene expression space where it is very unlikely to find cells, which would require to have thousands of cells just for a small network, or to take into account explicitly the value of time in the inference method. The latter point would require to have analytical results on the temporal distribution of cells in the gene expression space, which is generally out of range for realistic models like (1). For example, using the temporal dynamics of the probability vector *µ*_*t*_ on the basins for reconstructing the transition rates (see Section 2.3) would be challenging as an analytical link between these transition rates and the GRN can only be obtained up to a prefactor, which does not depend on the scaling factor *ε* but is specific to each transition [51]. The only solution seems then to combine analytical results with simulations, in the same philosophy than [5], but this leads to much more complex and time-consuming algorithms that the one which is developed in this article. To our point of view, this also argues against methods where a mixture model is used for reconstructing the temporal dynamics of a metastable process from stationary distributions [37].

### 6.3 Applicability of CARDAMOM to real single-cell datasets

The algorithm CARDAMOM has been shown to reconstruct accurately a most-likely GRN from timestamped *in silico* datasets, through the estimation of the metastable parameters associated to these data. It is then natural to ask whether the method could deal with real datasets or not. The method for estimating the metastable parameters should be well suitable for real datasets, provided that the Negative Binomial mixture approximation of mRNAs is accurate, and we have seen in Section 5.6 that the algorithm is suitable for realistic number of genes and cells. The main difficulty that could appear would then be related to the decrease in the performance of the algorithm with the number of modes appearing in the sample. Indeed, we recall that the regression step aims to infer a GRN matrix of size *n × n* from the values of the functions *k*_*on,i*_ at each attractor. First, it implies that too few clusters in the sample would mean too little information for the inference. Second, as developed in Section 5.5, too many clusters for each gene would make the sigmoidal bursts rate functions unsuitable, which could also lower the performances of the algorithm. Then, the level of multistability seems then to be critical for the accuracy of the inference. For the first point, we argue that regarding the noisy nature of mRNA counts, it is impossible to infer a reliable GRN when there is not enough multistability in the sample, *i*.*e* when the marginal distribution of mRNA associated to each gene is described by a simple negative binomial, the parameters of which does not evolve in time. We discuss in Section 6.4 the implications of the second point in terms of modeling.

To go further, we recall that the mathematical analysis beyond CARDAMOM lays on the point of view that cell differentiation processes can be coarsely reduced into a discrete process on a limited number of cell types, *i*.*e* that the main ingredients for characterizing stochasticity are the frequency modes describing the cell types and the random transitions between them, which is commonly accepted since [18]. Such transitions have been for example recently proposed as the basis for facilitating the concomitant maintenance of transcriptional plasticity and stem cell robustness [55]: in this case, the authors had proposed a phenomenological view of the transition dynamics between states, and our work may typically connect this cellular plasticity to an underlying GRN dynamics. This connection between a GRN and associated cell states should also be used for the quantitative modelling of stochastic state transitions underlying the generation of diversity in cancer cells [56], [14], or for investigating transitions between the large diversity of clusters that can be observed in human ES cell differentiation datasets [29].

### 6.4 Gene expression as a neural network

Interestingly, CARDAMOM appears close to a machine learning approach. Indeed, since each function *k*_*on,i*_ is independent from the others, depending only on the *i*^*th*^ line of the matrix *θ*, the regression step is fairly close to parallel regressions of each gene on the others. The choice of sigmoidal bursts rate functions makes these regressions appear as the learning step of an artificial neural network, which is simply a set of one-layer perceptron for each gene coupled by the crossed-penalization between the symmetric coefficients of the matrix *θ* (see (15)). In the light of this framework, the first step of the algorithm can be seen as an identification of the outputs from the data, which corresponds to the modes associated to each mRNA count. For sigmoidal bursts rate functions, we have seen in Section 5.5 that it is reasonable to consider only two modes. This makes the clustering step close to the prepossessing of a Boolean network analysis, which consists in assigning every cell to a state of a Boolean model, where a 0 may rather take any values between 0 and 1. Without crossed-penalization, the second step would exactly consist in learning the parameters of a perceptron as represented in Figure 13, where each line of the matrix *θ* corresponds to the weight of the perceptron. The method then builds an interesting link between machine learning and dynamical modeling.

**Figure 13:**
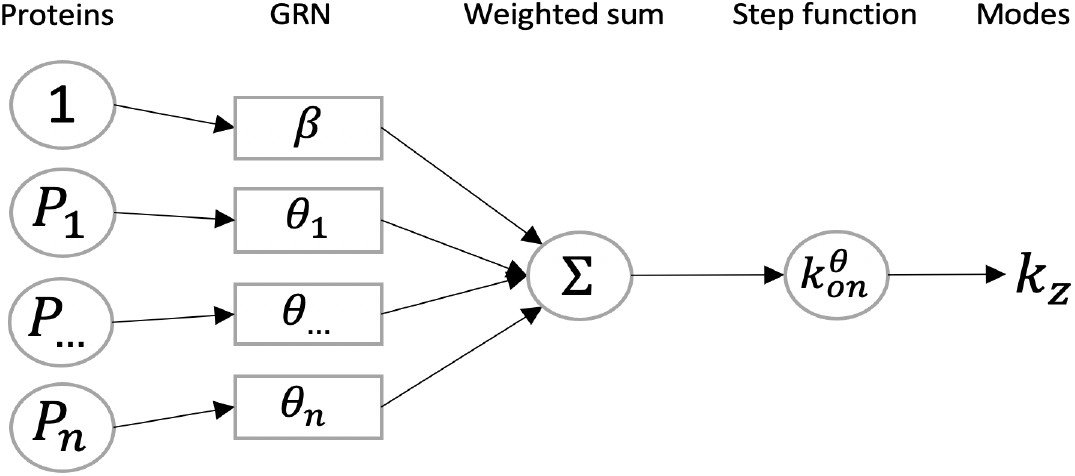
CARDAMOM can be interpreted as the learning of a perceptron, where the clustering step (Section 4.2.1) corresponds to the classification of the data, regarding to their associated modes of frequency, and the regression step (Section 4.2.2) to the identification of the weights of the perceptron, corresponding to the GRN.

The results presented in Section 5.5 let us suppose that the sigmoidal bursts rate functions of the form (2) have a limited complexity, *i*.*e* that they are not able to catch a multistability which would be associated to more than two potential wells for each gene. Thus, when applying the clustering step to a dataset showing an important multimodality, at least for some genes, the sigmoidal function may be not adapted for modeling the behaviour of the underlying process.The clustering step of CARDAMOM, described in Section 4.2.1 could therefore be seen as a preliminary to the choice of the “right” bursts rate function that should be used for minimizing the risk (17) (see Section 4.2.2): this function should be chosen in accordance with the number of modes detected for each gene. The neural network framework introduced above is interesting, as it suggests that the choice of the bursts rate function could correspond to the choice of the number of layers in the neural network of Figure 13. In that point of view, the work achieved in this paper treats the basic case of one layer, which could be extended, although the interpretability of the nodes for more than one layer remains an open question. We believe that this adaptability is necessary when dealing with highly complex single-cell data.

## Conclusion

We proposed in this work an efficient method for performing GRN inference from single-cell data, seen as the calibration of a mechanistic model of gene expression involving mRNA and protein levels. To the best of our knowledge, our approach is the first one which uses explicitly the popular notion of approximate developmental landscape, through the concept of metastability, for performing the inference. The method relies on a previous analysis of the mechanistic model developed in [51]. It provides a modular algorithm which consists in two steps: in a first time characterizing the metastable parameters associated to the coarse-grained model which describes the best the data, and in a second time solving a serie of regression problems aiming to link these parameters to a most-likely GRN. The method is implemented as a Python package called CARDAMOM, which is available on open-access. This algorithm seems to be very accurate for small but complex networks, and its computational speed allows to make it suitable for realistic number of genes and cells. Furthermore, such inference method based on a mechanistic model has the great advantage to make quantitative predictions that can then be easily tested by simulating the model. The simplicity of the regression step is particularly remarkable, and appears close to the learning step of a neural network. We believe that in addition to efficiently infer GRNs from timestamped datasets while keeping a high interpretability, CARDAMOM,which combines explicitly machine learning methods with mathematical modeling, could pave the way for new adaptive methods.

The next step would be to conceive bigger and more complex networks for which all the non-zeros entries of the GRN matrix do have an impact on the data when simulating the model, which is in itself a challenge, and to compare the performances of CARDAMOM to state-of-the-art algorithms used for GRN inference. As the problem of inference from a mechanistic model like (1) has been recently studied elsewhere using distinct mathematical tools [15], a collaboration is in progress for realizing an important benchmark of our method together with another algorithm specifically developed for facing transcriptional bursting and well-known methods such as GENIE 3 [19] and PIDC [8], and studying the ability of the mechanistic model to reproduce *in vitro* expression datasets when the model is calibrated by CARDAMOM.

## Appendix A Description of the parameters used for the model

We provide here a list of the main variables that are used throughout the article.

1. For the gene expression model:
  - *θ* a matrix defining the interactions between genes, corresponding to a matrix with diagonal terms defining external stimuli,
  - *k*_0,*i*_ is the basal rate of expression of gene *i*,
  - *k*_1,*i*_ is the maximal rate of expression of gene *i*,
  - *β*_*i*_ is the basal activity of gene *i*, which can be also considered as the constant activity of set of genes which are not measured and act on the network,
  - *d*_0,*i*_ is the degradation rates for mRNAs of gene *i*,
  - *d*_1,*i*_ is the degradation rate for proteins of gene *i*,
  - *s*_0,*i*_ is the creation rate for mRNAs of gene *i*,
  - *s*_1,*i*_ is the creation rate for proteins of gene *i*,
  - *k*_*off* ,*i*_ is the exponential rate of switching from state *on* to state *off* for the promoter of gene *i*,
  - *k*_*on,i*_ is a sigmoidal function depending on the global protein field *P ∈* Ω, defined in (2), which characterizes the exponential rate of switching from state *off* to state *on* for the promoter of gene *i*,
  - 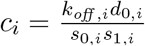 is the exponential rate of proteins of gene *i* that are created at every burst, in the model (3).
2. For the numerical analysis allowing to understand the algorithm CARDAMOM:
  - *Z* denotes a set of basins of attraction associated to the deterministic limit for a given GRN, and (*P*_*z*_)_*z∈Z*_ the set of corresponding attractors,
  - *k*_*z,i*_ = *k*_*on,i*_(*P*_*z*_) corresponds to frequency mode of the promoter of gene *i* associated to the basin *z ∈ Z*,
  - *µ*_*t*_ is a probability vector describing the probability for a cell to be in each basin *z ∈ Z* at time *t. μ*denotes this probability vector at the steady state,
  - 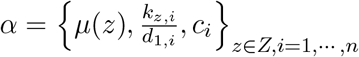 is the set of parameters which describe a Gamma mixture of the form (6),
  - 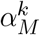 is the set of parameters which describe a Gamma mixture associated to a gene expression matrix 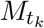 measured at time *t*_*k*_,
  - 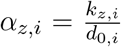 denotes the renormalized frequency mode for the promoter of gene *i* of a cell within a basin *z ∈ Z*, that is accessible from scRNA-seq data.

## Appendix B Master equation of the reduced model and explicit stationary distributions

The master equation on the probability density *u*(*t, ·*) of the bursty model (3), describing only proteins, associated to a GRN *θ* appears as an integro-differential equation:

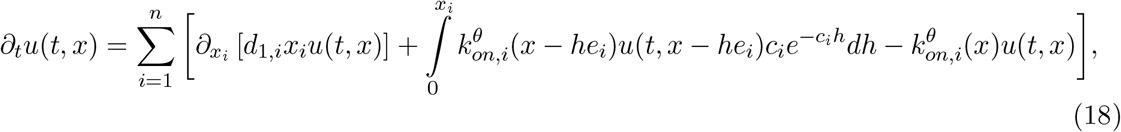

where each *e*_*i*_ is a vector of size *n* with only zero entries except on the *i*^*th*^ position.

## Appendix C Regression method in case of proteomic data

In this section, we show how CARDAMOM, which aims to infer a GRN from scRNA-seq data, can be interpreted as a restriction of a more general algorithm where mRNAs are seen as a proxy for the proteins levels, which uses more intensively the characteristics of the Gamma mixture approximation (6) instead of the modes only.

In the model (1), a GRN is completely encoded through the functions 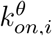. Then, it is straight- forward that knowing the exact values taken by these functions on the whole protein space would allow to determine the value of the associated GRN. The method described in Section 3.1 consists in using the value of these functions on the set of the attractors, but we did not directly use the information provided by the probability vector on the basins *µ*. Given a experimentally observed distribution of proteins, it would be fruitful if the mixture approximation allowed to evaluate the functions *k*_*on,i*_ not only on the attractors of the basins but also on the position of the cells that are observed. For this sake, we define the functions 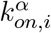 for all *i* = 1, *…, n*, for all *x ∈* Ω:

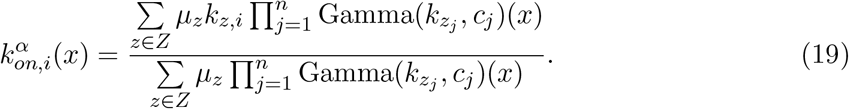

The two following lemmas are proved at the end of this section:

### Lemma 1.

*Replacing* 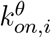 *by* 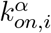 *in the model* (3), *the stationary distribution is exactly given by the mixture distribution* (6).

### Lemma 2.

*We consider the evolution of a population of cells, initially distributed under a Gamma mixture û*_*α*_ *of the form* (6), *in the model* (3) *driven by the functions* 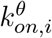. *We have the inequality:*

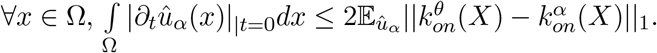

Lemma 1 suggests that the difference between the distributions *û*_*θ*_ and *û*_*α*_, which is supposed to be small according the the analysis of Section 2, should be related to the difference between the two classes of parametric functions 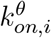 and 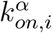. The inequality of Lemma 2 gives a more precise reason for considering this difference. Indeed, the expected value of the 1-norm of the difference between these two functions is an upper bound of a quantity which measures how rapidly the associated mixture distribution is going to change when it is taken as an initial condition in the master equation of the model (3) driven by 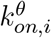, that we call the impulsion of this mixture. We remark that for every product of Gamma distributions centered on one of the attractors associated to a GRN, the expected value on the right hand side of the inequality of Lemma 2 is expected to be small, as the cells are not going to jump to another basin in short times. The “right” mixture associated to a GRN *θ* is then the one which is going to be accurate in the highest part of the gene expression space, *i*.*e* that takes into account the highest number of basins. But a simple sum of Gamma mixture would not be accurate on the areas of the gene expression space where the Gamma distributions cross each other, as it does not take into account the potential depth associated to each attractor. The balance *µ*(*z*) can then be seen as a balance which characterizes the most-likely energetic barrier between the potential wells for reconstructing the right steady-state behavior. Note that this ability of the vector *μ* to identify the depth of the basins should be closely linked to the ability of a quasipotential to describe the transitions between the basins [51].

We compare in Figure 14 the functions 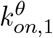 and 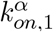 for the toggle-switch network described in Table 1, illustrating the fact that the approximation seems accurate inside the basins and allows to reproduce quite accurately the basins of attraction of the deterministic system, although the functions on the boundary of the basins have a slightly different behaviour. This is in line with the ideas developed in Section 4.2.3.

**Figure 14:**
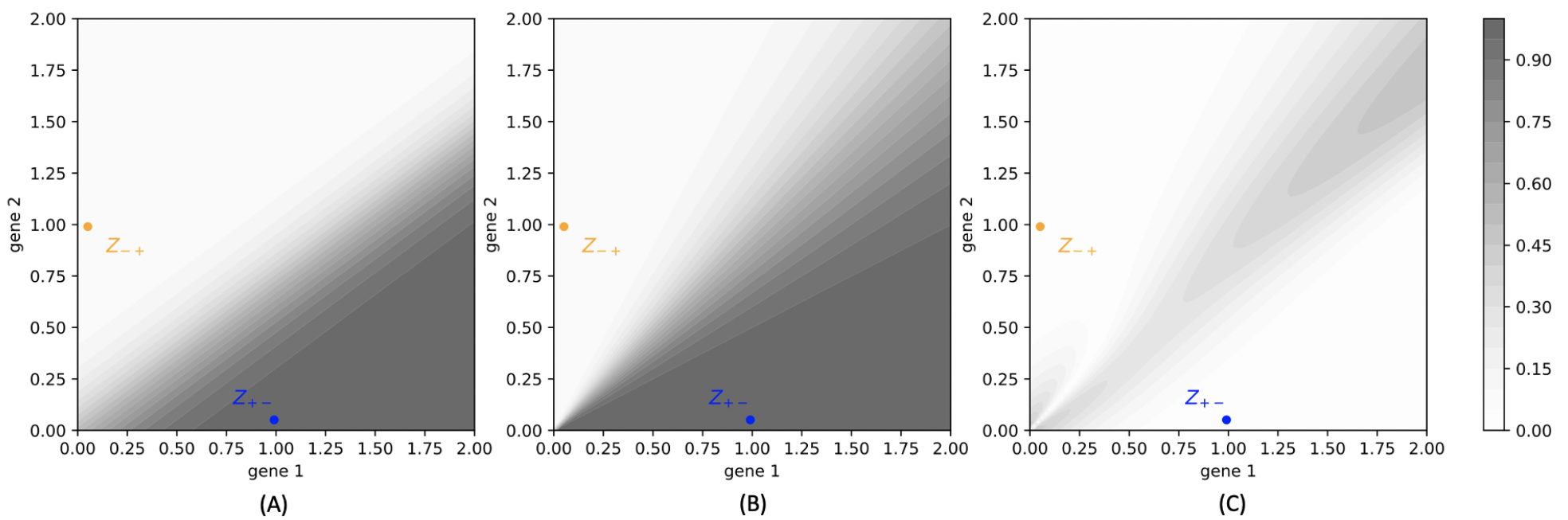
(A): Color map of the function 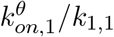 characterized by the toggle-switch described in Table 1, on the gene expression space. (B): Color map of the function 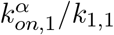 characterized by the mixture parameters associated to the same network, obtained with the method described in Figure 4, on the gene expression space. (C): Color map of the function 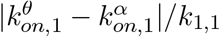, on the gene expression space.

The upper bound of the impulsion of a mixture distribution in the model driven by a GRN *θ*, which is provided by the inequality of Lemma (2), appears as a good choice for the function *R* described in Section 3.2. It is natural to substitute the distribution *û*_*α*(*X*)_ by the empirical distribution associated to the matrix *X*. We also decide to take the 2-norm in order to simplify the minimization problem. We would finally obtain instead of (17):

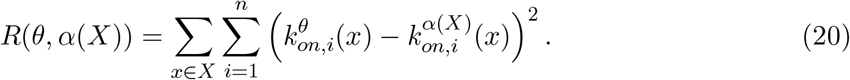

We remark that the function to be optimized in CARDAMOM (17) corresponds exactly to this new function (20) when the cells *x* are exactly located on the attractors. This is in line with the fact that in our framework, the most likely position for the proteins vector *P* knowing mRNAs are on the attractors of the basins to which the cell belongs. This lack of information cannot be compensated: indeed, it would either suppose that an observed deviation of a count of mRNA from an attractor is better explained by the fact that the proteins are far from their most-likely position than mRNA itself, which is not the case since mRNAs are known to be much more noisy than proteins, either that the functions *k*_*on,i*_ are far from their value on the attractors even when proteins are far from the boundary, in which case the Gamma mixture approximation (6) would not be accurate and our method is no more relevant.

### Proof of Lemma 1

Injecting the functions 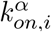 instead of 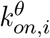 and the distribution *û*^*α*^ of the form (6) instead of *u* on the right-hand side of the master equation (18), we obtain for all *i* = 1, *…, n*:

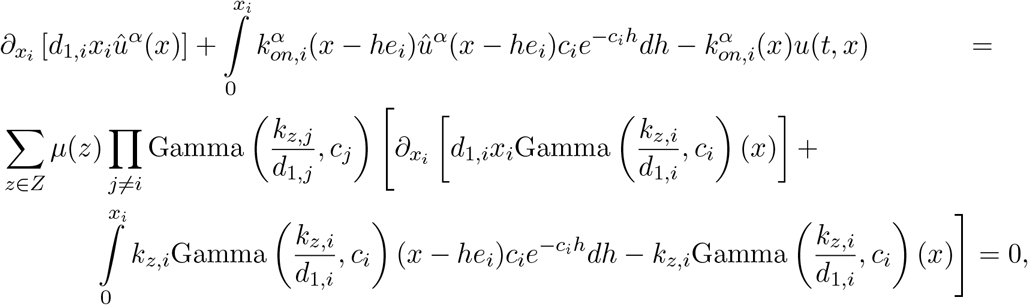

because Gamma 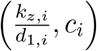 is the unique stationary distribution of the model (3) for one gene and a constant *k*_*on,i*_ = *k*_*z,i*_ [24]. Then, the right-hand side of the master equation is null and *û*^*α*^ is the unique stationary distribution of the model (3) when the burst rates functions are of the form (19).

### Proof of Lemma 2

This lemma follows from the previous Lemma 1. Indeed, from the master equation at *t* = 0, defining *u*(0, *x*) = *û*^*α*^(*x*), we have:

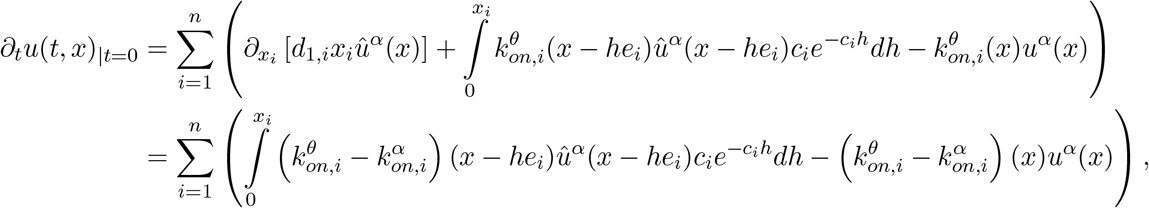

where the second equality comes from the fact that *uû*^*α*^ is the stationary solution of the master equation (18) with burst rates functions 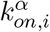. Thus, we obtain:

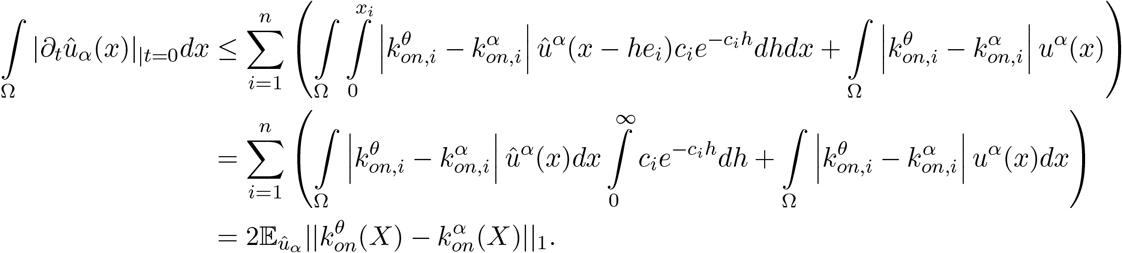

## Code availability

The algorithm CARDAMOM presented in Section 4.2, as well as the code for generating the figure 7, are available on https://gitbio.ens-lyon.fr/eventr01/cardamom.

The algorithm used for simulating the model as well as to generate the tree-like networks used in Section 5.6 can be found on https://github.com/ulysseherbach/harissa.

## Acknowledgment

This work was supported by funding from French agency ANR (SingleStatOmics; ANR-18-CE45-0023-03). I would like to thank especially Thibault Espinasse for enlightening discussion and statistical advices, as well as Ulysse Herbach for having highlighted the notion of bursts modes for the model of gene expression, Thomas Lepoutre and Olivier Gandrillon for critical reading of the manuscript. I also thank all members of the SBDM and Dracula teams, and of the SingleStatOmics project, for providing such stimulating working environment. I finally thank the BioSyL Federation, the LabEx Ecofect (ANR-11-LABX-0048) and the LabEx Milyon of the University of Lyon for inspiring scientific events.

